# Rank Deficiency in Parameter Estimation for Hodgkin–Huxley Type Current Models

**DOI:** 10.64898/2026.11.16.767153

**Authors:** Matthias Heinkenschloss, Saina Namazifard, Fabrizio Gabbiani

## Abstract

The Jacobians of nonlinear least squares residuals play a central role when simultaneously fitting parameters of Hodgkin–Huxley (HH) type models to ionic currents measured through voltage clamp experiments. Their rank deficiency is tied to parameter degeneracy, i.e., the ability to produce the same current response with different parameters, and the Jacobians’ null spaces provide insight into the structure of the parameter degeneracy, as well as into which additional experiments can reduce or even remove it. This paper analyzes Jacobian rank structure across HH-type models used for H-, A-type, fast sodium, and delayed rectifier currents. The Jacobians are rank deficient to varying degrees, depending on experimental conditions. Analytical representations of the Jacobians’ null spaces are provided. In all models, one source of rank deficiency is the multiplication of the peak conductance and state variables, such as activation and inactivation, inherent to the HH-type model structure. H-current models derived from either whole cell, or cell-attached patch recordings exhibit additional rank deficiencies, implying the local existence of manifolds of parameters that lead to the same current responses. For A-type, fast sodium, and delayed rectifier models there is no additional rank deficiency, although their Jacobians may exhibit characteristics associated with ill-conditioned parameter estimation problems. Descriptions of the null spaces of these Jacobians are used to prove that the rank deficiency can be reduced or even be removed by fixing certain parameters, or by adding additional experiments. The analytical results are illustrated numerically, using data from voltage clamp experiments described in the literature.

## 1 Introduction

Since the application of cable theory to single neurons (Rall 1962), their biophysics has been described by compartmental models endowed with ion channels of Hodgkin–Huxley (HH) type (Hodgkin and Huxley 1952). Over the past several decades, multiple methods have been devised to efficiently fit such models to experimental data (Druckmann et al. 2007; Huys et al. 2006; Keren et al. 2005), a topic that remains the subject of active research (Deistler et al. 2025). It is now also well established that models may exhibit parameter degeneracy in the sense that different combinations of model parameters generate virtually identical neuron responses (Prinz et al. 2004; Achard and De Schutter 2006). Understanding parameter degeneracy in compartmental neuron models is an important topic since it provides a unique way to assess how robust neuronal processing is and what type of constraints are imposed on single neuron computation by its biophysical components (Goaillard and Marder 2021). The Jacobians of the residuals in the nonlinear least squares problems used to estimate the parameters, specifically the ranks of these Jacobians, play a critical role in understanding this parameter degeneracy.

In this paper, we compute the ranks of Jacobians associated with several voltage clamp experiments covering the range of experimentally relevant biophysical ion channel models. While the importance of parameter degeneracy in single neuron models has long been recognized and studied through numerical simulations, little attention has been devoted to understanding the mathematical structure of their associated solution manifolds. Because the rank properties of Jacobians and their null spaces are critical to the existence of local solution manifolds and to their computation, this paper provides theoretical foundations crucial to understanding parameter solution manifolds. We focus on a simple case: the possibility of parameter degeneracy in HH-type ion channel models fitted to voltage clamp data. This setting is advantageous because the fitting problem is simplified by the membrane potential being held constant. As a result, the time-evolution of the gating functions under voltage clamp conditions for the studied ion channel can be expressed in closed form after solving analytically the associated first order differential equations.

In a nutshell, parameters in voltage clamp experiments are estimated as solutions of a nonlinear least squares problem

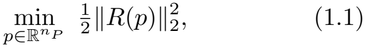

where the parameter vector, 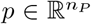, consists of all model parameters fitted simultaneously such as the ion channel peak conductance, reversal potential, gating variable(s) steady-state values and time constants. The components of the residual function are given by the differences between the current *I*(*t*_*j*_; *V*_*C,i*_; *p*) at time *t*_*j*_ generated with the voltage clamp potential *V*_*C,i*_ and the model parameters *p*, and the corresponding measured current *I*_meas_(*t*_*j*_; *V*_*C,i*_) from experiments. That is, the residual function has components

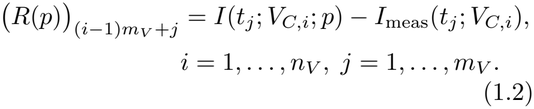

The residual vector 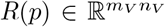 with components (1.2) has *n*_*V*_ blocks, each corresponding to one voltage clamp potential, *V*_*C,i*_, and containing the difference between the predicted current and the measured current at *m*_*V*_ sampled time points, *t*_*j*_. Associated with the residual function 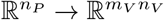 is its Jacobian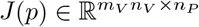, the matrix of partial derivatives of *R*. For *k* = (*i* − 1)*m*_*V*_ + *j, i* = 1, …, *n*_*V*_, *j* = 1, …, *m*_*V*_, and *l* = 1, …, *n*_*P*_, the (*k, l*)-th entry of the Jacobian is *J*_*kl*_(*p*) = *∂R*_*k*_(*p*)*/∂p*_*l*_ = *∂I*(*t*_*j*_; *V*_*C,i*_; *p*)*/∂p*_*l*_.

The Jacobian is an important ingredient in the characterization of solutions to nonlinear least squares problems and to their numerical solution (e.g., Björck 2024, Chap. 8; Dennis and Schnabel 1996, Chap. 10; Nocedal and Wright 2006, Chap. 10). It also plays a central role in assessing the quality of the estimated parameters, e.g., through the estimation of parameter confidence regions and other statistical properties (e.g., Bard 1974; Donaldson and Schnabel 1987; Seber and Wild 2003). Finally, the Jacobian is critically involved in determining parameter degeneracy, as we will describe next.

Assume that *p*_*\**_ solves the nonlinear least squares problem (1.1). Moreover, suppose that locally around *p*_*\**_ there exists a smooth manifold, i.e., a smooth map 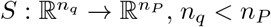, such that *S*(*q*_*\**_) = *p*_*\**_ and *R*(*S*(*q*)) = *R*(*p*_*\**_) for all *q* in a neighborhood of *q*_*\**_. Then 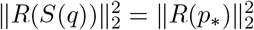 and all *p* = *S*(*q*) for *q* in a neighborhood of *q*_*\**_ also solve the nonlinear least squares problem (1.1). The existence of such a map *S* can be proven under the assumptions of the so-called constant rank theorem. See Fracasso and Heinkenschloss (2027) and, in slightly different form, Eriksson et al. (2005). Moreover, the columns of the Jacobian 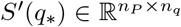 span the null space of the Jacobian *J* (*p*_*\**_) (Fracasso and Heinkenschloss 2027). Thus, knowledge of the Jacobian column rank and its null space is critical in determining the existence of parameter degeneracy and in computing the local solution manifold, respectively (Namazifard et al. 2026, Section 4.2.4; Fracasso and Heinkenschloss 2027). To illustrate these points, consider the following simple geometrical example.

Let the parameter vector be two dimensional, *p* = (*p*_1_, *p*_2_)^*T*^, and given *β*_1_, *β*_2_ *>* 0, consider the function

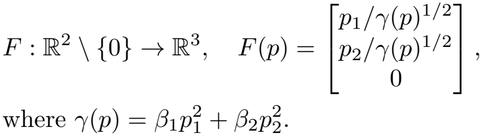

This function maps rays emanating from 0 *∈* ℝ ^2^ onto an ellipse in the two-dimensional subspace of ℝ^3^ spanned by *e*_1_ = (1, 0, 0)^*T*^ and *e*_2_ = (0, 1, 0)^*T*^ . Now, consider minimizing the squared norm of the residual *R*(*p*) = *F* (*p*) *− z*, where *z ∈* ℝ^3^, *z* ≠ 0. Geometrically, this corresponds to finding the point, *x*, on the ellipse defined by *F* closest to *z*, see Fig. 1. The Jacobian of *R* is the Jacobian of *F* and is given by

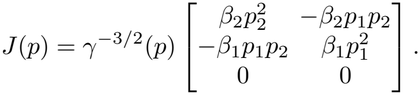

**Fig. 1.**
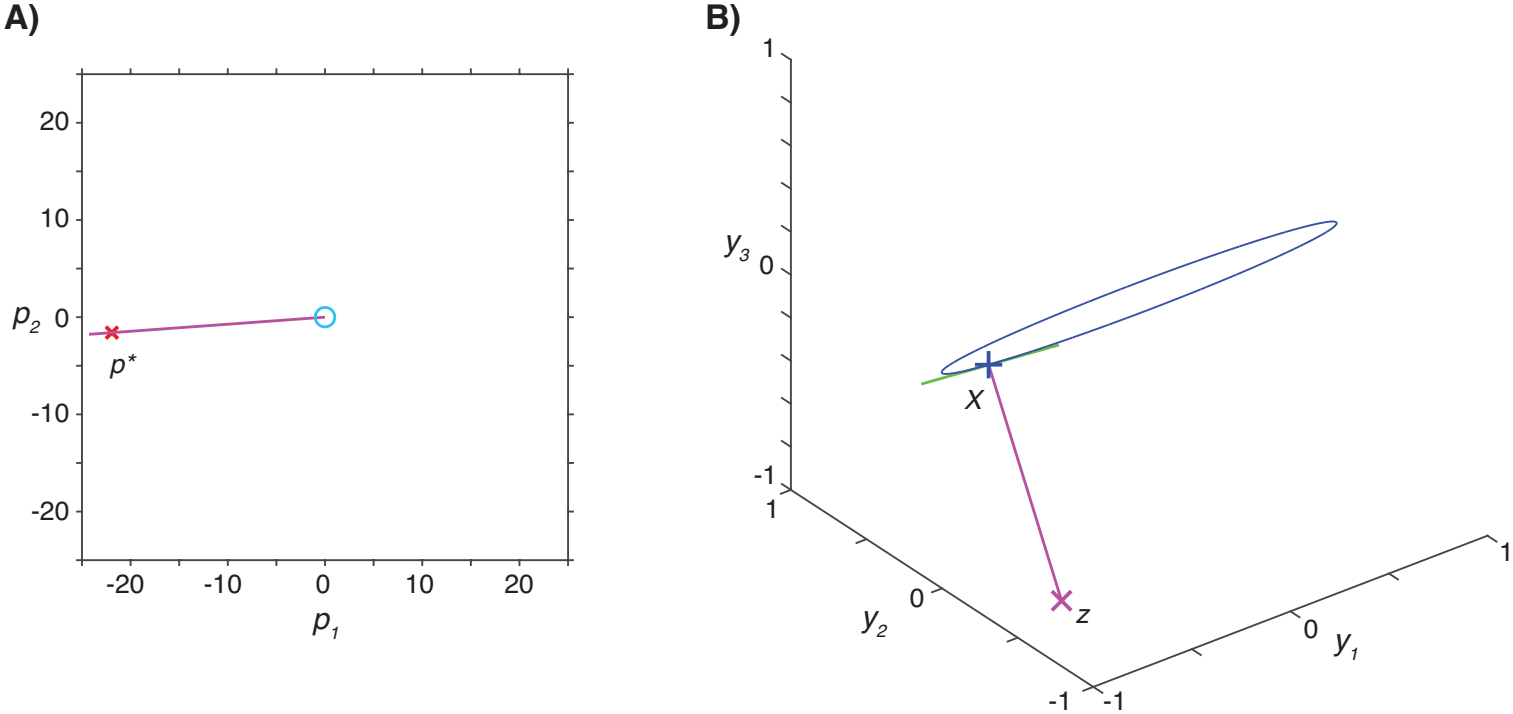
Geometrical example of solution manifold: **(A)** In parameter space *p*_*\**_ (red cross) lies on the ray originating at 0 (magenta line), which is the solution manifold mapped onto *x* by *F* . The cyan line is the unit circle. **(B)** The ellipse onto which ℝ^2^*\ {*0*}* is projected (blue line), and the point *x* = *F* (*p*_*\**_) (blue plus sign) which is the point on the ellipse closest to *z* (magenta cross). The green line is the tangent vector at *x* to the ellipse, which is perpendicular to the magenta line joining *x* and *z*.

For all *p ≠* 0, *J* (*p*) has rank one, and the null space of *J* (*p*) is spanned by *p* = (*p*_1_, *p*_2_)^*T*^ . If *p*_*\**_ solves the nonlinear least squares problem, then any 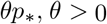, also solves the nonlinear least squares problem.

As mentioned earlier, we compute the ranks of the Jacobians associated with several voltage clamp experiments covering the range of experimentally relevant biophysical ion channel models. Additionally, in the rank deficient case, we give explicit formulas for a basis of the null space of the Jacobian, which determine the local geometry of the solution manifold. In a related paper (Namazifard et al. 2026), we use the results derived here to develop methods that efficiently trace the associated parameter solution manifold away from a local solution *p*_*\**_. In another related extension, the same results allow one to determine joint parameter correlations and confidence regions in non-degenerate cases (Namazifard et al. 2026).

The rest of this paper is organized as follows. We first pose the general parameter estimation problem for voltage clamp experiments in Section 2.1. In Section 2.2, we prove that the Jacobian associated with parameter estimation of the simplest HH-type model, used to describe the H-current present in many neurons (He et al. 2014), is rank deficient by two, once parameter redundancies due to the formulation of the model are accounted for. Thus, voltage clamp data alone defines a two-dimensional solution manifold for the HH-type H-current model. We then show that adding two further experiments each alleviates the rank deficiency by one, yielding a full rank Jacobian. In Section 2.3, we consider the related problem of constraining the H-current model through cell-attached patch clamp recordings. The rank deficiency is at most equal to one but surprisingly depends on the holding potential selected for the experiments. In this case, we show that one additional experiment is sufficient to yield a full rank Jacobian. In Section 2.4, we show that an HH-type model of an inactivating current, such as the A-type current, has no rank deficiency once the parameter redundancies due to the formulation of the model are accounted for. In Section 2.5, we extend these results to the Jacobians associated with the two currents used in the HH model of action potential generation. Proofs of the results in Sections 2.2 and 2.3, and partial proofs of the results in Sections 2.4 and 2.5 were derived by hand computations. The algebraic complexity of the current models in Sections 2.4 and 2.5 prompted us to use artificial intelligence to suggest complete proofs of the results in these sections, which we then modified for clarity and consistency. Finally, Section 3 numerically illustrates the results by computing the singular value spectrum of the models considered in Sections 2.2– 2.5 for experimentally fitted currents extracted from the literature.

## 2 Parameter Estimation in Voltage Clamp Experiments

### 2.1 Hodgkin–Huxley-Type Compartmental Neural Models

The Hodgkin–Huxley (HH) formalism, originally introduced to describe action potential generation in the squid giant axon (Hodgkin and Huxley 1952), provides a general framework for modeling neuronal membrane dynamics. The general form of an HH-type model is

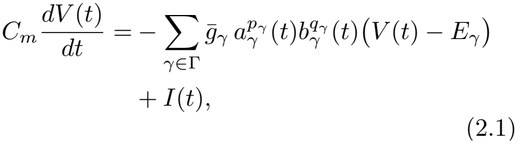

where *C*_*m*_ is the membrane capacitance, and Γ is a set of *n*_Γ_ channels. Further, 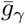 and *E*_*γ*_ are the maximal conductance and reversal potential of ionic current *γ*, respectively, and *a*_*γ*_ and *b*_*γ*_ are voltage-dependent activation and inactivation gating functions, respectively, with corresponding integer exponents *p*_*γ*_ and *q*_*γ*_ taking values ranging from 1 to 4 for activation, and 0 or 1 for inactivation. The gating functions *a*_*γ*_, *b*_*γ*_, *γ ∈* Γ, are modeled by differential equations, which for a generic gating function *y ∈ {a*_*γ*_, *b*_*γ*_ : *γ ∈* Γ*}* is given by

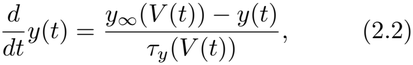

where *y*_*∞*_(*V* ) *∈* [0, 1] denotes the voltagedependent steady-state (in-) activation and *τ*_*y*_(*V* ) *>* 0 is the corresponding (in-) activation time constant.

We investigate parameter estimation in HH-type models under voltage clamp conditions. During a voltage clamp experiment, the membrane potential is held at a prescribed holding potential (*V*_*C*,hold_) then stepped instantaneously to a constant command voltage (*V*_*C*_) at time *t*_*V*,0_, and held at this constant command voltage during the time [*t*_*V,0*_, *t*_*V,f*_ ). Therefore, for *t ∈* [*t*_*V,0*_, *t*_*V,f*_ ), the rate of membrane voltage change with respect to time is zero (d*V/*d*t* = 0), and (2.1) reduces to

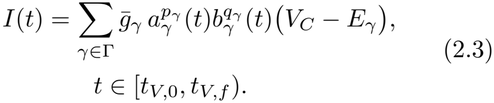

Because *V* (*t*) = *V*_*C*_ is constant for *t ∈* [*t*_*V,0*_, *t*_*V,f*_ ),, (2.2) reduces to the linear differential equation

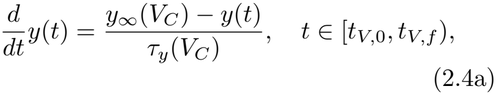

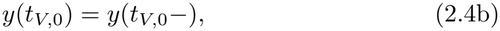

where *y*(*t*_*V*,0_*−*) is the value of the gating function before the membrane potential is stepped to *V*_*C*_. Using the method of the integrating factor, the solution of (2.4) is computed as

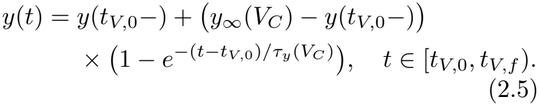

Note that *y*(*t*) *→y*_*∞*_(*V*_*C*_) as *t →∞* . In a voltage clamp experiment, it is assumed that the membrane potential is held at a prescribed holding potential (*V*_*C*,hold_) for long enough that *y*(*t*_*V*,0_*−*) can be replaced by *y*_*∞*_(*V*_*C*,hold_), so that the solution of (2.4) becomes

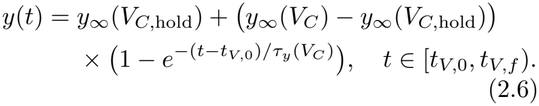

Inserting (2.6) for *y∈ {a*_*γ*_, *b*_*γ*_ : *γ ∈* Γ *}* into (2.3) gives

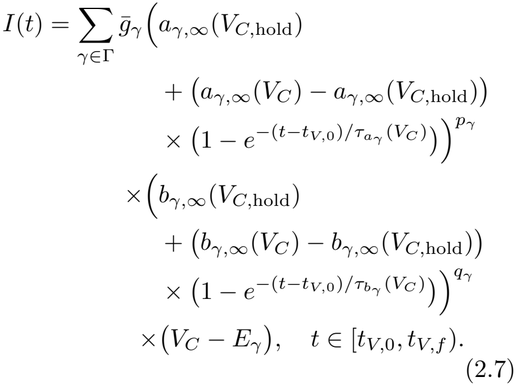

Under experimental voltage clamp conditions, the number of ion channels is either *n*_Γ_ = 2 or 1, corresponding to the ‘whole cell’ and ‘cell-attached’ voltage clamp configurations, respectively (Hamill et al. 1981). In the first case, the measured current comprises that of the studied ion channel, *S*, usually isolated pharmacologically, and the resistive leak current of the membrane, *L*, so that Γ = *{L, S}* . In the second case, the studied ion channel is isolated from a small patch of membrane and no leak current is present, Γ = *{S}* . The leak does not have any associated gating variables, *p*_*L*_ = *q*_*L*_ = 0, and is modeled through its conductance 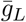 and reversal potential *E*_*L*_. The studied current *S* always has an activation variable (*p*_*S*_ *>* 0). Thus, the number, 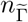, of gating variables is either 1 or 2, depending on whether *q*_*S*_ = 0 or 1.

The voltage clamp experiment is repeated with *n*_*V*_ different command voltages 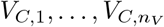. The goal in the whole cell voltage clamp experimental configuration is to estimate

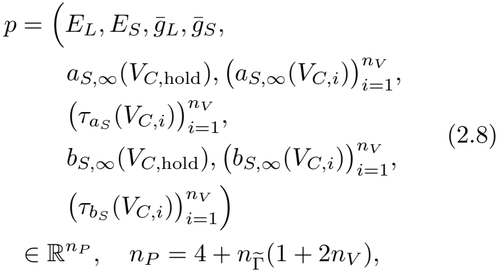

from the model (2.7) with *V*_*C*_ replaced by *V*_*C,i*_ and corresponding current measurements *I*_meas_(*t*_*V,j*_ ; *V*_*C, i*_) at times *t*_*V,j*_ *∈* [*t*_*V,0*_, *t*_*V,f*_ *), j* = 1, …,*m*_*V*_. In the cell-attached configuration the number of parameters is 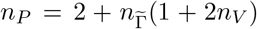 and since there is no leak current (i.e.,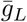and *E*_*L*_ are removed from *p* in (2.8)). We determine estimates for the parameter vector as the solution of the nonlinear least squares problem

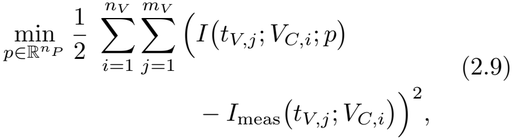

where 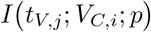 is the current model (2.7) with *V*_*C*_ replaced by *V*_*C,i*_ at time 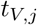 and 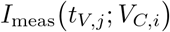 is the corresponding current measurement. Abstractly, (2.9) can be written as (1.1), where 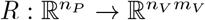 is the residual function with components (1.2).

### 2.2 hyperpolarization-Activated Ionic Current – Whole Cell Voltage Clamp

First, we consider a voltage clamp experiment for a current governed by a single activation variable with an exponent *p*_*H*_ = 1. A typical example is the ubiquitous hyperpolarization-activated current (H-current) found in many neurons (He et al. 2014). In the Numerics Section 3, we consider as an example the H-current of the lateral pyloric (LP) neuron in the stomatogastric ganglion (STG) of crabs (Golowasch and Marder 1992; Liu et al. 1998).

#### 2.2.1 Voltage clamp experiment

The general HH-type formulation introduced in Section 2.1 is used to model a voltage clamp experiment on currents such as the H-current. The model is of the form (2.7) with Γ = *{L, H}, p*_*L*_ = *q*_*L*_ = 0, *p*_*H*_ = 1, *q*_*H*_ = 0, and *a*_*H*_ = *h*,

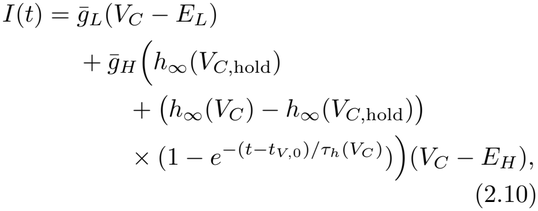

for *t ∈* [*t*_*V,0*_, *t*_*V,f*_ ),where 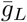 and 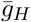 are the maximal conductances of the leak and H-current, respectively, and *E*_*L*_ and *E*_*H*_ are their corresponding reversal potentials.

For *n*_*V*_ voltage clamp recordings, we estimate the parameters

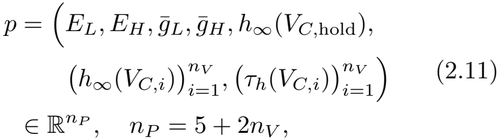

as a solution of the nonlinear least squares problem (2.9), where *I (t*_*V,j*_ ; *V*_*C,i*_; *p*) is the function (2.10) with *V*_*C*_ = *V* _*C,i*_ at times 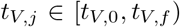, and *I*_meas_ *(t*_*V,j*_ ; *V*_*C*,_ *i)* are the measured currents. For a numerical example, see Section 3 and Fig. 2A.

**Fig. 2.**
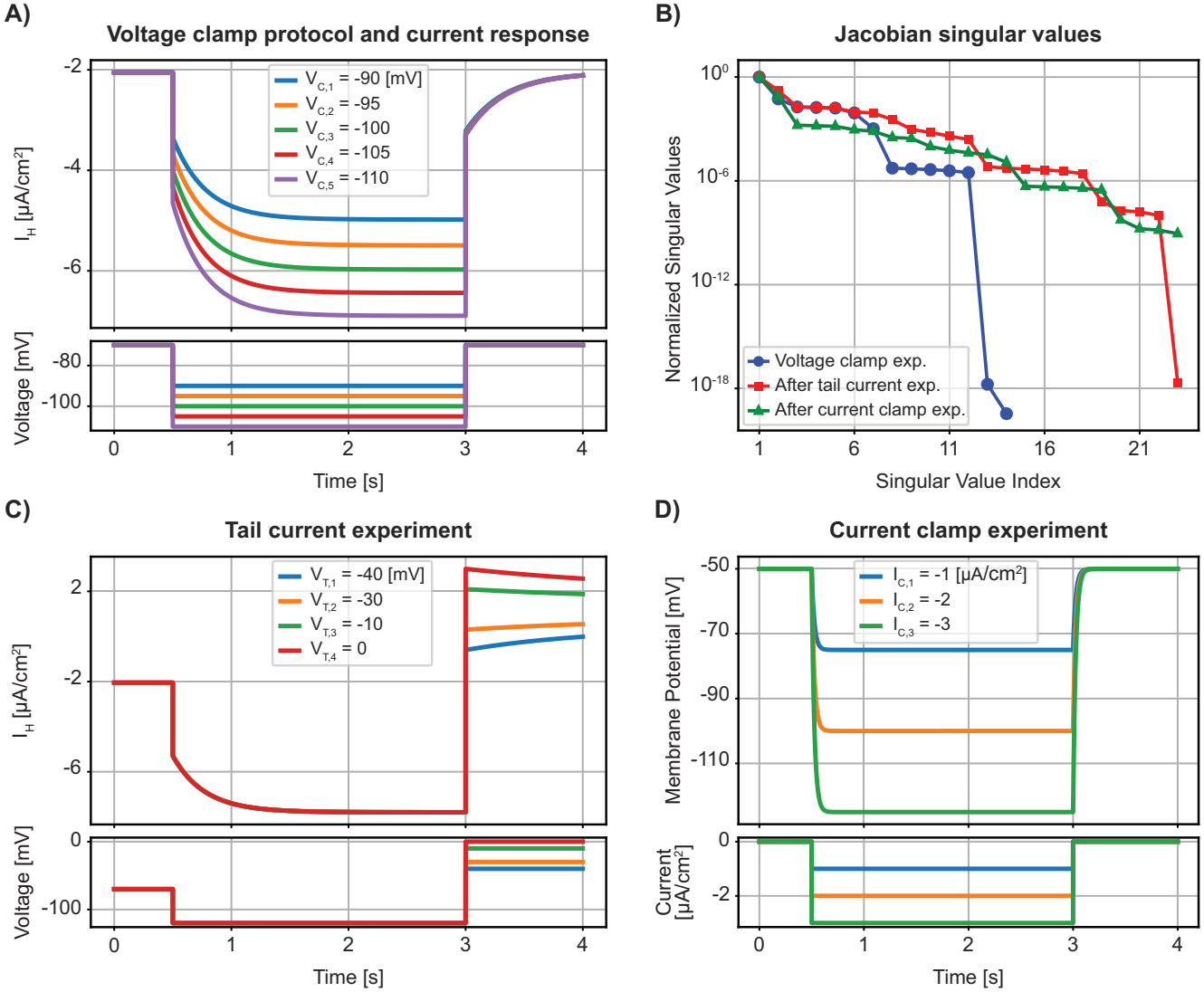
Experimental protocols and normalized singular values for the STG LP H-current model: **(A)** Voltage clamp traces for the H-current model. **(B)** Normalized singular values of the Jacobian evaluated at the corresponding model parameters. **(C)** Tail current traces used to decrease the Jacobian rank deficiency by one. **(D)** Current clamp traces used to decrease the Jacobian rank deficiency by one.

Although we have much more data than parameters to identify, *n*_*V*_ *m*_*V*_ *≫ n*_*P*_, the Jacobian 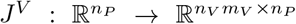 of the residual function corresponding to (2.9) is rank deficient, and its rank is independent of parameters (2.11) satisfying the following biophysically meaningful assumptions.

##### Assumption 2.1.

*The command voltages satisfy* 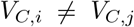 *for i ≠ j (if n*_*V*_ = 1, *this condition is void), and the parameters satisfy* 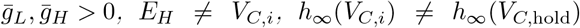 *and τ*_*h*_*(V*_*C*_,_*i*_*) > 0, i = 1*, …, *n*_*V*_

##### Theorem 2.2.

*Let the command voltages and parameters* (2.11) *satisfy Assumption 2*.*1, and let* 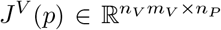 *be the Jacobian corresponding to* (2.9).

1. *If n*_*V*_ *>* 1 *and m*_*V*_ *>* 2, *then the Jacobian J*^*V*^ (*p*) *has rank n*_*P*_ *−* 3 = 2 + 2*n*_*V*_.
2. *If n*_*V*_ = 1 *and m*_*V*_ *>* 2, *then the Jacobian J*^*V*^ (*p*) *has rank n*_*P*_ *−* 4 = 3.

*Proof* The proof is given in Appendix A.1.

In (2.10), 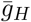 and *h*_*∞*_(*V*_*C*,hold_), *h*_*∞*_(*V*_*C,i*_) appear as products. Therefore, we cannot identify 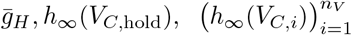 simultaneously. One of these parameters needs to be fixed and removed from the estimation problem to resolve this model-based degeneracy. In the proof of Theorem 2.2 in Appendix A.1 it is shown that the vector

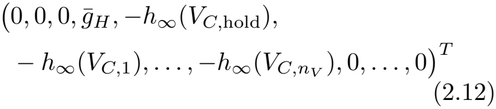

is in the null space of the Jacobian of the residual function corresponding to (2.9). For example, (2.12) shows that if 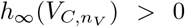, the (5 + *n*_*V*_ )-th column corresponding to the partial derivative with respect to 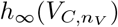 in the Jacobian can be expressed using the other columns. The null space vector (2.12) reflects the fact that 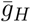 appears only in a product with 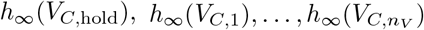 in (2.10) and, therefore, these parameters can never be identified simultaneously.

Under judicious selection of the voltage clamp protocol, the H-current will be fully activated at the most hyperpolarized voltage step, which we assume is 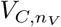. Thus, we fix

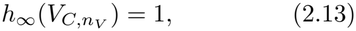

and estimate the parameters

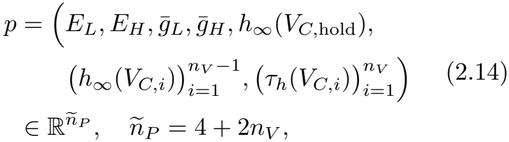

by minimizing (2.9) while fixing (2.13). In this case, the rank deficiency of the associated Jacobian reduces by one.

##### Corollary 2.3.

*Let the command voltages and parameters* (2.14) *satisfy Assumption 2.1, and let* 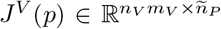 *be the Jacobian corresponding to* (2.9).

i. *If n*_*V*_ *>* 1 *and m*_*V*_ *>* 2, *then the Jacobian J*^*V*^ (*p*) *has rank ñ*_*P*_ *−* 2 = 2 + 2*n*_*V*_ .
ii. *If n* = 1 *and m >* 2, *then the Jacobian J*^*V*^ (*p*) *has rank ñ*_*P*_ *−* 3 = 3.

*Proof* The result follows from Theorem 2.2 and the fact that by (2.12) the column of the Jacobian of (2.9) corresponding to the partial derivative with respect to 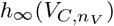 can be expressed as a linear combination of the other columns.

The main result, part *i*. of Corollary 2.3, is illustrated numerically in Section 3 (Fig. 2B, blue line and data points).

#### 2.2.2 Tail current experiment

To decrease the rank deficiency of the Jacobian *J*^*V*^ (*p*), we add a so-called tail current experiment to the previous voltage clamp experiment. In the tail current experiment, the membrane potential is first held at a large hyperpolarizing voltage, *V*_*T*,hold_, to activate the H-current, and then *n*_*T*_ voltage steps that bracket the expected value of *E*_*H*_ are applied, see Fig. 2C. The measured current at the *n*_*T*_ selected voltage steps is called the tail current. The unknown parameters associated with the tail current experiment are the steady-state activation at the holding potential, *h*_*∞*_(*V*_*T*,hold_), the steady-state activation values at the *n*_*T*_ tail voltage steps, *h*_*∞*_(*V*_*T,i*_), and the corresponding activation time constants, *τ*_*h*_(*V*_*T,i*_). Therefore, including the tail current data introduces 1 + 2*n*_*T*_ parameters to the estimation problem. Consequently, when *n*_*T*_ tail current recordings are added to the previous *n*_*V*_ voltage clamp recordings, we need to estimate the parameters

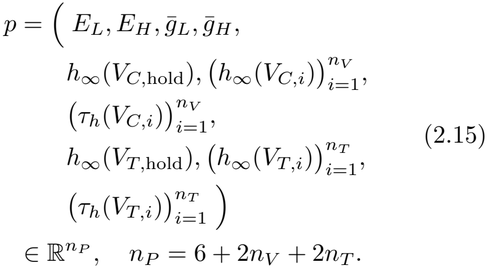

The parameters are estimated by solving the least squares problem

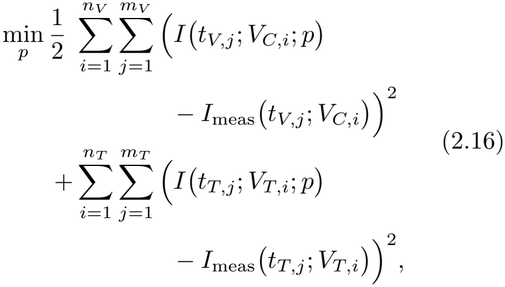

where the terms in the first double sum are given as before, *I (t*_*T,j*_ ; *V*_*T,i*_; *p*) is the function (2.10) with *V*_*C*,hold_, *V*_*C*_ replaced by *V*_*T*,hold_, *V*_*T,i*_, and at *t*_*V,0*_ = *t*_*T,0*_ times *t*_*T,j*_ ∈ *[t*_*T,0*_, *t*_*T,f*_ ), and *I*_meas_ (*t*_*T,j*_ ; *V*_*T,I*_ ) are the measured currents.

The nonlinear least squares problem (2.16) is of the form (1.1), where 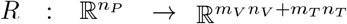 is the function with components given by the expressions within the squared terms in (2.16). We can compute the Jacobian 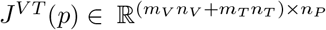 of this residual function and analyze its rank. Theorem 2.5 below shows that adding the tail current experiment reduces the rank deficiency, but the Jacobian *J*^*V T*^ (*p*) is still rank deficient, and its rank is independent of the parameter choice.

##### Assumption 2.4.

*Assumption 2.1 holds. Furthermore, the command voltages satisfy* 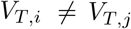 *for i ≠ j (if n*_*T*_ = 1, *this condition is void), and the parameters satisfy V*_*T,i*_ *≠ E*_*H*_, *h*_*∞*_(*V*_*T,i*_) *≠ h*_*∞*_(*V*_*T*,hold_), *h*_*∞*_(*V*_*C*,hold_) ≠ *h*_*∞*_(*V*_*T*,hold_), *and τ*_*h*_(*V*_*T,i*_) *>* 0, *i* = 1, …, *n*_*T*_ .

##### Theorem 2.5.

*Let the command voltages and parameters* (2.15) *satisfy Assumption 2*.*4, let n*_*V*_ *>* 1 *and m*_*V*_ *>* 2, *and let* 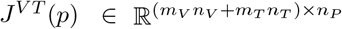 *be the Jacobian corresponding to* (2.16).

i. *If n*_*T*_ *>* 1 *and m*_*T*_ *>* 2, *then the Jacobian J*^*V T*^ (*p*) *has rank n*_*P*_ *−* 2 = 4 + 2*n*_*V*_ + 2*n*_*T*_ .
ii. *If n*_*T*_ = 1 *and m*_*T*_ *>* 2, *then the Jacobian J*^*V T*^ (*p*) *has rank n*_*P*_ *−* 3 = 5 + 2*n*_*V*_ .

*Proof* The proof is given in Appendix A.2.

As shown in the proof of Theorem 2.5 in Appendix A.2, the first column vector in (A19) is the vector in the null space of the Jacobian *J*^*V T*^ (*p*) that reflects the fact that 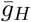 appears only in a product with *h*_*∞*_ (*V*_*C*,hold_), 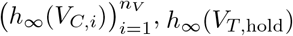, and 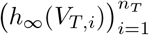 in (2.10) and, therefore, these parameters can never be identified simultaneously. If 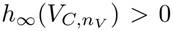, the (5 + *n*_*V*_ )- th column corresponding to the partial derivative with respect to 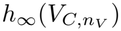 in the Jacobian (A13) can be expressed using the other columns. If we again select 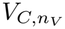 such that the channel is fully open, fix 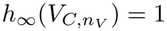 (see (2.13)), and estimate the remaining parameters

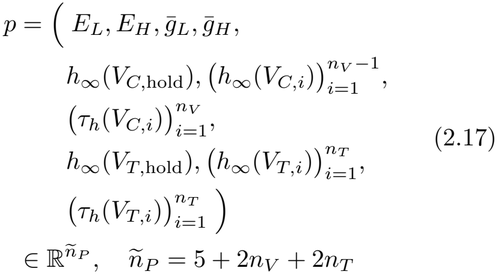

by solving the corresponding problem (2.16), the rank deficiency of the associated Jacobian reduces by one. This leads to the following corollary of Theorem 2.5.

##### Corollary 2.6.

*Let the command voltages and parameters* (2.17) *satisfy Assumption 2*.*4, let n*_*V*_ *>* 1 *and m*_*V*_ *>* 2, *and let* 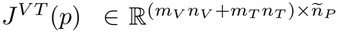 *be the Jacobian corresponding to* (2.16) *with* 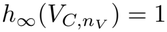.

i. *If n*_*T*_ *>* 1 *and m*_*T*_ *>* 2, *then the Jacobian J*^*V T*^ (*p*) *has rank ñ*_*P*_ *−* 1 = 4 + 2*n*_*V*_ + 2*n*_*T*_ .
ii. *If ñ*_*T*_ = 1 *and m*_*T*_ *>* 2, *then the Jacobian J*^*V T*^ (*p*) *has rank ñ*_*P*_ *−* 2 = 5 + 2*n*_*V*_ .

*Proof* The result follows from Theorem 2.5 and the fact that the column of the Jacobian of (2.16) corresponding to the partial derivative with respect to 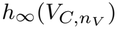 can be expressed as a linear combination of the other columns.

The main result, part *i*. of Corollary 2.6, is illustrated numerically in Section 3 (Fig. 2B, red line and data points).

Comparing Theorems 2.2 and 2.5 shows that if *n*_*V*_ *>* 1 voltage clamp recordings are performed and *n*_*T*_ *>* 1 tail current recordings are added, the rank deficiency of the Jacobian reduces by one.

#### 2.2.3 Current clamp experiment

To eliminate the remaining rank deficiency, we consider adding a current-clamp experiment, assuming that the H-current has been pharmacologically blocked, yielding 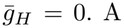 constant current injection, *I*_*C*_, is then initiated at time *t*_*C*,0_ and continues until *t*_*C,f*_ . Following (2.1), the membrane voltage dynamics are described by

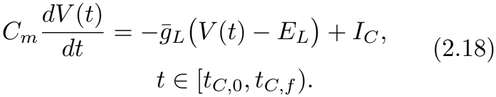

Using the method of integrating factors, the solution of (2.18) is

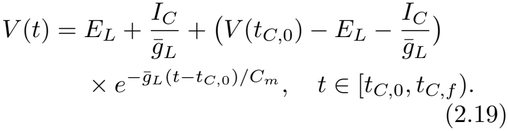

In the interval [0, *t*_*C*,0_) no current is injected (*I*_*C*_ = 0) and the voltage remains at the initial condition *V* (0) = *E*_*L*_. As a result, *V* (*t*_*C*,0_) = *E*_*L*_. Then in *n*_*C*_ recordings, constant current injections 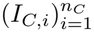 are applied and the corresponding voltage is measured at times *t*_*C,j*_ ∈ [*t*_*C,0*_, *t*_*C,f*_), *j* = 1, …, *m*_*C*_. The membrane capacitance *C*_*m*_ is assumed to be known. A numerical example is treated in Section 3 (Fig. 2D).

Unlike the tail current experiment, the current clamp experiment does not introduce additional unknown parameters. As a result, when *n*_*V*_ voltage clamp recordings, *n*_*T*_ tail current recordings, and *n*_*C*_ current clamp recordings are used simultaneously, we still estimate the parameters in (2.15) or those remaining after fixing 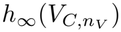 but now as the solution of

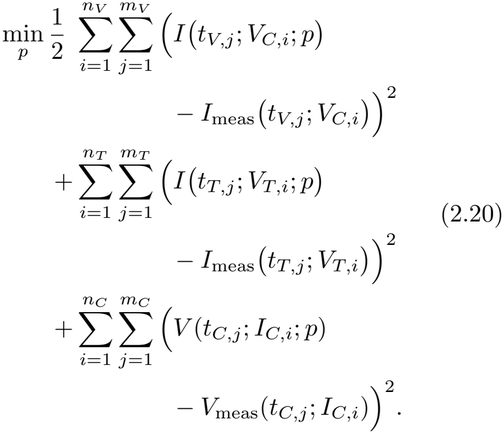

In the third double sum, *V* (*t*_*C,j*_ ; *I*_*C,i*_; *p*) is computed from (2.19) with *V* (*t*_*C*,0_) = *E*_*L*_, *I*_*C*_ = *I*_*C,i*_, *i* = 1, …, *n*_*C*_, and *V*_meas_(*t*_*C,j*_; *I*_*C,i*_), *j* = 1, …, *m*_*C*_, are the measured voltages.

The nonlinear least squares problem (2.20) is again of the form (1.1), where 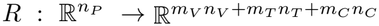 is the function with components given by the expressions within the squared terms in (2.20) and the parameters are given by (2.15) or (2.17). The Jacobian associated with this least squares problem is denoted by 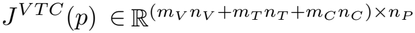 . The next results show that the addition of current clamp recordings results in a Jacobian with a null space of dimension one if the parameters (2.15) are used, and a full column-rank Jacobian if 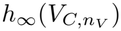 is fixed.

##### Assumption 2.7.

*Assumption 2*.*4 holds. Furthermore, let n*_*V*_ *>* 1 *and m*_*V*_ *>* 2, *and let either (a) n*_*C*_ *≥* 2 *and I*_*C,i*_ *≠I*_*C,j*_ *for i ≠ j, or (b) m*_*C*_ *≥* 2 *and* 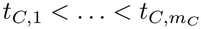.

The following theorem considers the *n*_*P*_ = 6 + 2*n*_*V*_ + 2*n*_*T*_ parameters (2.15).

##### Theorem 2.8.

*Let the voltage clamp, current clamp and model parameters* (2.15) *satisfy Assumption 2*.*7 and let* 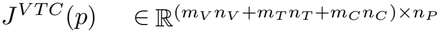 *be the Jacobian corresponding to* (2.20).

i. *If n*_*T*_ *>* 1 *and m*_*T*_ *>* 2, *then the Jacobian J*^*V T C*^(*p*) *has rank n*_*P*_ *−* 1 = 5 + 2*n*_*V*_ + 2*n*_*T*_.
ii. *If n*_*T*_ = 1 *and m*_*T*_ *>* 2, *then the Jacobian J*^*V T C*^(*p*) *has rank n*_*P*_ *−* 1 = 7 + 2*n*_*V*_ .

*Proof* The proof is given in Appendix A.3.

If 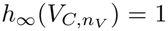 is fixed (see (2.13)) and the *ñ*_*P*_ = *n*_*P*_ *−* 1 = 5+2*n*_*V*_ +2*n*_*T*_ parameters (2.17) are used, the corresponding Jacobian has full column rank.

##### Corollary 2.9.

*Let the voltage clamp, current clamp and model parameters* (2.17) *satisfy Assumption 2*.*7, then the Jacobian* 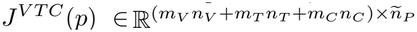 *has full column rank, ñ*_*P*_.

*Proof* The result follows from Theorem 2.8 and the fact that the column of the Jacobian of (2.20) corresponding to the partial derivative with respect to 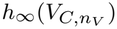 can be expressed as a linear combination of the other columns.

This result is illustrated numerically in Section 3 (Fig. 2B, green line and data points).

### 2.3 Hyperpolarization-Activated Ionic Current – Cell-Attached Voltage Clamp

Isolation of an ionic current can affect the Jacobian rank of the associated HH-type model. For instance, in Angelo et al. (2007), the cell-attached voltage clamp configuration applied to Purkinje cells of the cerebellum eliminates the presence of a leak current, yielding a pure H-current. Therefore, by removing the leak current in (2.10), the current model is

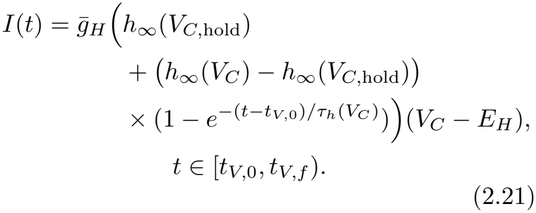

#### 2.3.1 Voltage clamp experiment

For *n*_*V*_ voltage clamp recordings, we estimate the parameters

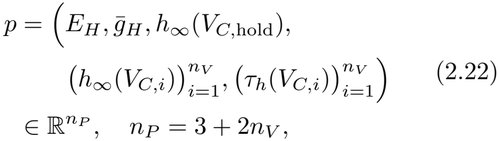

as the solution of the nonlinear least squares problem (2.9) where *I* (*t*_*V,j*_; *V*_*C,i*_; *p*) is the function (2.21) with *V*_*C*_ = *V*_*C,i*_ at times *t*_*V,j*_ *∈* [*t*_*V*,0_, *t*_*V,f*_ ), and *I*_meas_ *(t*_*T,j*_ ; *V*_*T,I*_ ) are the measured currents. For a numerical example, see Section 3 and Fig. 3A.

**Fig. 3.**
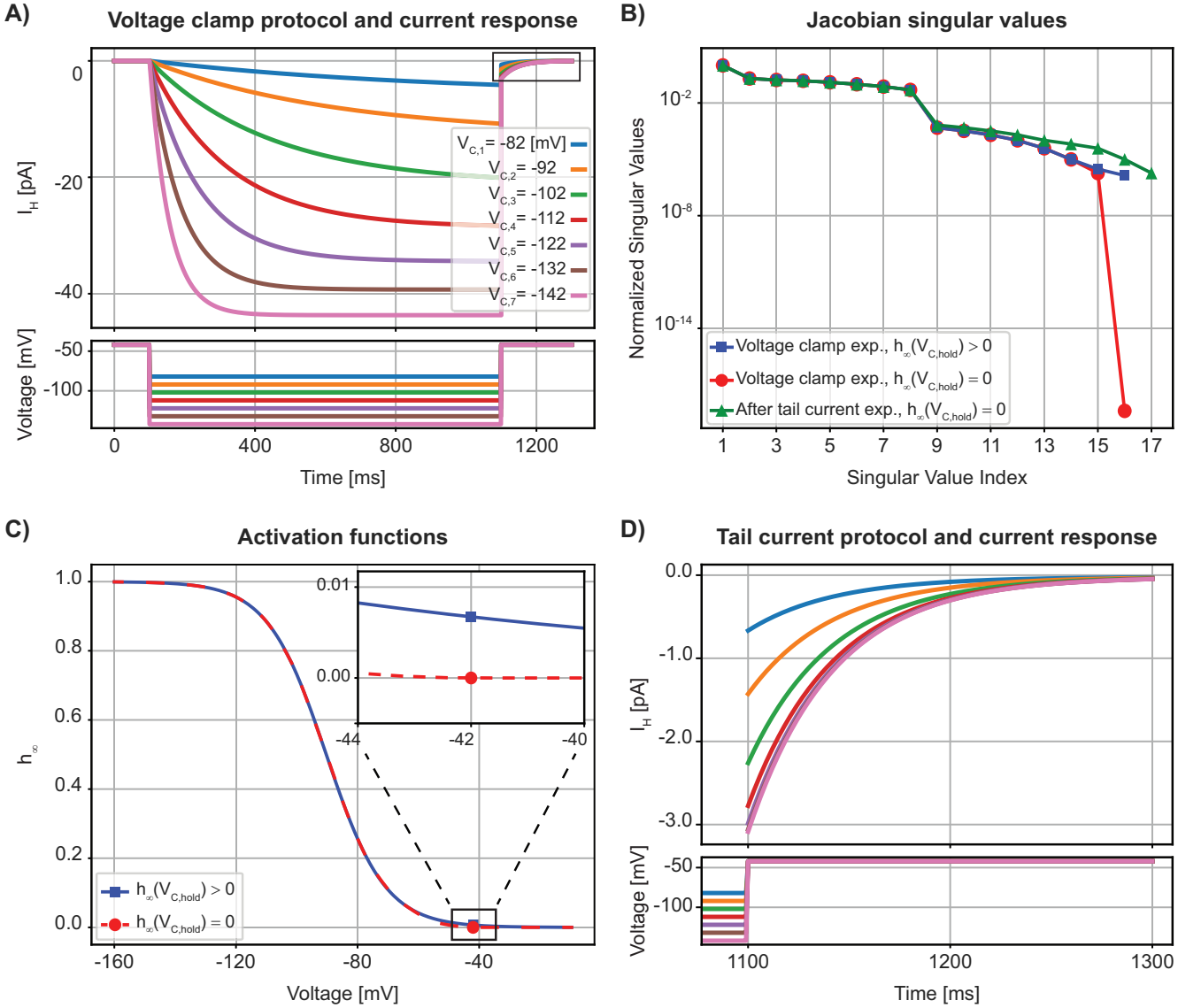
Experimental protocol, activation curve, and normalized singular values for the cell-attached Purkinje H-current model. **(A)** Voltage clamp traces for the dendritic cell-attached H-current model. **(B)** Activation function *h*_*∞*_ of the H-current model, with or without *h*_*∞*_(*V*_*C*,hold_) = 0 (red and blue lines and dots, respectively; the top right inset magnifies the two curves close to 0, and matches the black box in the main plot). **(C)** Normalized singular values of the Jacobian evaluated at the corresponding model parameters for the two activation functions shown in (B), with and without the tail current experiment shown in (D). **(D)** Tail portion of the voltage clamp recordings, marked by a black box in (A), top right.

It turns out that the removal of the leak channel substantially changes the rank properties of the Jacobian corresponding to (2.9). Moreover, its rank depends on whether *h*_*∞*_(*V*_*C*,hold_) = 0 or *h*_*∞*_(*V*_*C*,hold_) *>* 0.

##### Theorem 2.10.

*Let the command voltages and parameters* (2.22) *satisfy Assumption 2*.*1, and let* 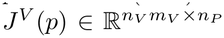 *be the Jacobian corresponding to* (2.9).

i *a. If n*_*V*_ *>* 1, *m*_*V*_ *>* 2, *and h*_*∞*_(*V*_*C*,hold_) = 0, *then the Jacobian J*^*V*^ (*p*) *has rank n*_*P*_ *−* 2 = 1 + 2*n*_*V*_ .
i *b. If n*_*V*_ *>* 1, *m*_*V*_ *>* 2, *and h*_*∞*_(*V*_*C*,hold_) *>* 0, *then the Jacobian J*^*V*^ (*p*) *has rank n*_*P*_ *−* 1 = 2 + 2*n*_*V*_.
ii *If n*_*V*_ = 1 *and m*_*V*_ *>* 2, *then the Jacobian J*^*V*^ (*p*) *has rank n*_*P*_ *−* 2 = 3.

*Proof* The proof is given in Appendix A.4.

As for the whole cell H-current LP neuron model considered in Section 2.2, we fix 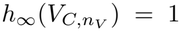 and remove it from the list of parameters to avoid model-based degeneracy due to the multiplicative combination of 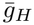 and *h*_*∞*_ in (2.21). The number of parameters to be identified is *ñ*_*P*_ = *n*_*P*_ *−* 1 = 2 + 2*n*_*V*_ .

##### Corollary 2.11.

*Let the command voltages and parameters* (2.22) *and* 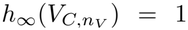 *satisfy Assumption 2*.*1, and let* 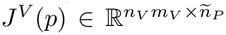*be the Jacobian corresponding to* (2.9).

i *a. If n*_*V*_ *>* 1, *m*_*V*_ *>* 2, *and h*_*∞*_(*V*_*C*,hold_) = 0, *then the Jacobian J*^*V*^ (*p*) *has rank ñ*_*P*_ *−* 1 = 1 + 2*n*_*V*_.
i *b. If n*_*V*_ *>* 1, *m*_*V*_ *>* 2, *and h*_*∞*_(*V*_*C*,hold_) *>* 0, *then the Jacobian J*^*V*^ (*p*) *has full column rank ñ*_*P*_ = 2 + 2*n*_*V*_.
ii *If n*_*V*_ = 1 *and m*_*V*_ *>* 2, *then the Jacobian J*^*V*^ (*p*) *has rank ñ*_*P*_ *−* 1 = 3.

*Proof* The proof follows from Theorem 2.10 and the fact that the column of the Jacobian of (2.9) corresponding to the partial derivative with respect to 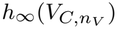 can be expressed as a linear combination of the other columns, as is shown in the proof of Theorem 2.10 in Appendix A.4.

The main result, part *i*. of Corollary 2.11, is illustrated using two numerical examples in Section 3 (Fig. 3C, red and blue lines and data points, respectively). The corresponding activation curves, with *h*_*∞*_(*V*_*C*,hold_) = 0 and *h*_*∞*_(*V*_*C*,hold_) *>* 0, respectively, are plotted in Fig. 3B.

Corollary 2.11 shows that even if we fix 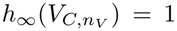, the Jacobian *J*^*V*^ (*p*) has a onedimensional null space if *h*_*∞*_(*V*_*C*,hold_) = 0 or if *n*_*V*_ = 1. Adding tail current recordings eliminates this rank deficiency, as we will show next.

#### 2.3.2 Tail current experiment

The tail current experiment we use here is modeled after, e.g., Angelo et al. (2007) and is different from that described in Section 2.2.2. Here, we use the same voltage clamp protocol and additionally fit the decaying tail portion of the current as the membrane potential is clamped back to its holding potential (*V*_*C*,hold_) from each command voltage (*V*_*C,i*_), see Fig. 3D. This introduces only one additional parameter to the problem, which is the activation time constant at the holding potential *τ*_*h*_(*V*_*C*,hold_). We therefore estimate

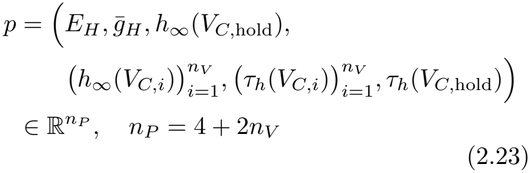

as the solution of the least squares problem

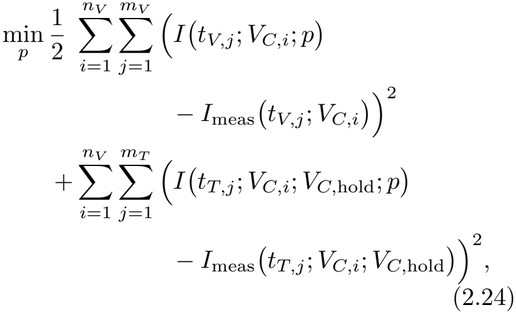

where the terms in the first double sum are given as before, *I* (*t*_*T,j*_; *V*_*C,i*_; *V*_*C*,hold_; *p*) is the tail current resulting from clamping the membrane potential back to its holding potential (*V*_*C*,hold_) from each command voltage (*V*_*C,i*_), and *I*_meas_ (*t*_*T,j*_; *V*_*C,i*_; *V*_*C*,hold_) are the corresponding measured tail currents. According to (2.6) with *y* = *h*, the gating function at *t*_*T*,0_, with *t*_*V*,0_ *< t*_*T*,0_ *< t*_*V,f*_, is

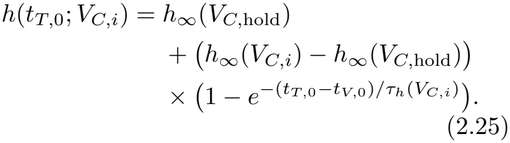

When the membrane potential is clamped back to its holding potential (*V*_*C*,hold_) the gating function *h* = *y* satisfies (2.4) with *t*_*T*,0_ = *t*_*V,f*_ and *h*(*t*_*T*,0_) = *h*(*t*_*V,f*_ *−*) from (2.25) and *V*_*C*_ = *V*_*C*,hold_. Thus, by (2.6),

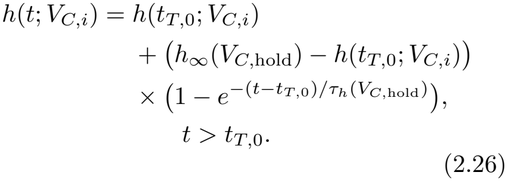

Inserting this into the current equation gives

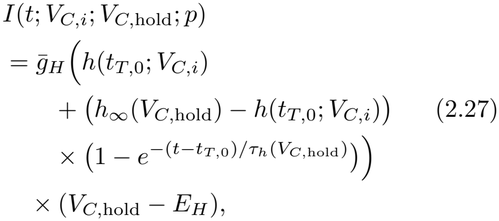

for *t ∈* [*t*_*T*,0_, *t*_*T,f*_ ), where *h*(*t*_*T*,0_; *V*_*C,i*_) is given by (2.25).

The following theorem shows that, after adding the tail current experiment, the Jacobian 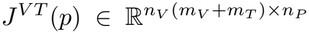 associated with the residual of the problem (2.24) has rank *n*_*P*_ *−* 1, for *h*_*∞*_(*V*_*C*,hold_) *≥* 0 and any *n*_*V*_ *>* 0.

##### Theorem 2.12.

*Let the command voltages and parameters* (2.23) *satisfy Assumption 2*.*1 and τ*_*h*_(*V*_*C*,hold_) *>* 0, *and let* 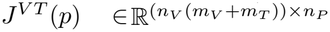 *be the Jacobian corresponding to* (2.24).

*If m*_*V*_ *>* 2 *and m*_*T*_ *>* 2, *then the Jacobian J*^*V T*^ (*p*) *has rank n*_*P*_ *−* 1 = 3 + 2*n*_*V*_ .

*Proof* The proof is given in Appendix A.5.

The proof of Theorem 2.12 shows that the vector in the null space of *J*^*V T*^ (*p*) corresponds to the parameter degeneracy due to the multiplicative combination of 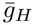 and *h*_*∞*_ in (2.21). Again, the degeneracy can be removed if we fix 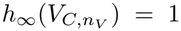 and remove it from the list of parameters. The number of parameters is now *ñ*_*P*_ = *n*_*P*_ *−* 1 = 3 + 2*n*_*V*_ .

##### Corollary 2.13.

*Let the command voltages and parameters* (2.23) *satisfy Assumption 2*.*1 and τ*_*h*_(*V*_*C*,hold_) *>* 0.

*If m*_*V*_ *>* 2 *and m*_*T*_ *>* 2, *then the Jacobian* 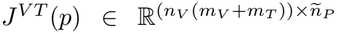 *corresponding to* (2.24) *with parameters* (2.23) *and* 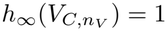 *has full column rank, ñ*_*P*_ = 3 + 2*n*_*V*_ .

*Proof* The proof follows from Theorem 2.12 and the fact that the column of the Jacobian of (2.24) corresponding to the partial derivative with respect to 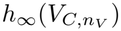 can be expressed as a linear combination of the other columns, as is shown in the proof of Theorem 2.12 in Appendix A.5.

This result is illustrated numerically in Section 3 (Fig. 3C, green line and data points).

### 2.4 A-Type Potassium Current

A-type potassium channels possess a fast activation and a slower inactivation gating function. Accordingly, the model follows the form presented in (2.1) – (2.7) with Γ = *{L, A}, p*_*L*_ = *q*_*L*_ = 0, *p*_*A*_ = 3, *q*_*A*_ = 1, *a*_*A*_ = *n*, and *b*_*A*_ = *h*. In the Numerics Section 3, we consider as an example the A-current of the STG LP neuron (Liu et al. 1998), see Fig. 4A.

**Fig. 4.**
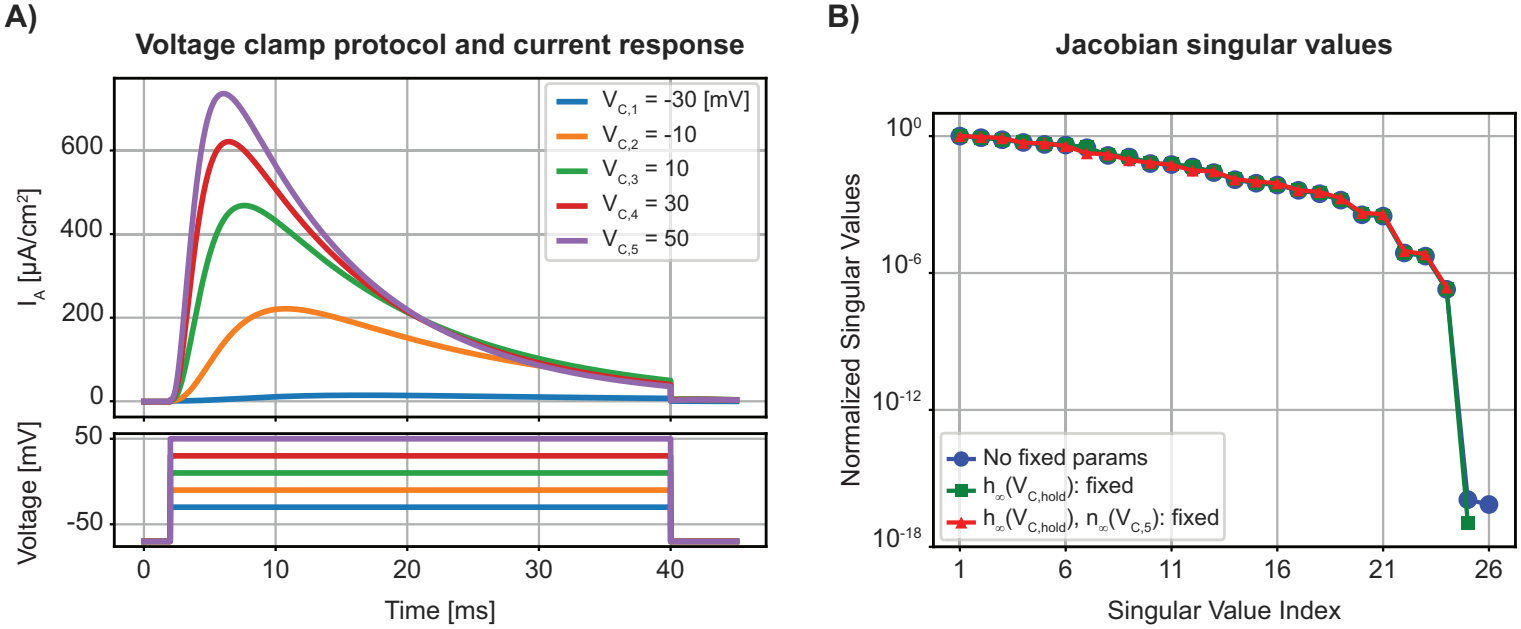
Parameter degeneracy in voltage clamp experiments for the STG A-current model: **(A)** Voltage clamp traces for the A-current model. **(B)** Normalized singular values of the Jacobian evaluated at the corresponding model parameters.

Similar to the analytical solution provided in (2.10), the current corresponding to a command voltage *V*_*C*_ can be written as follows

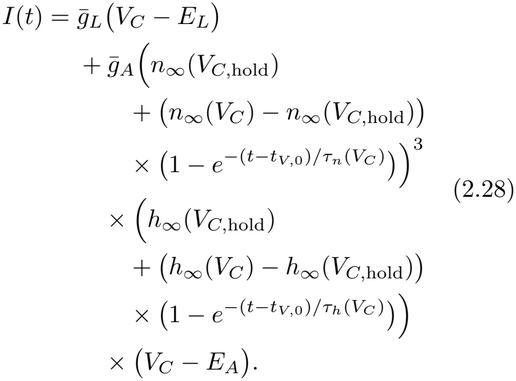

We have *n*_*V*_ voltage clamp recordings with command voltages *V*_*C,i*_, *i* = 1, …, *n*_*V*_ . We estimate the unknown parameters

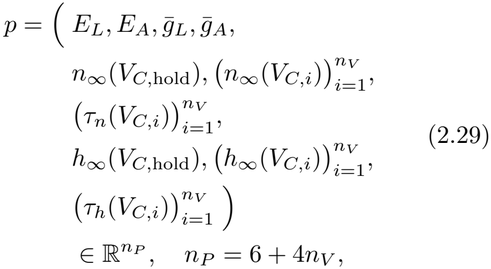

as the solution of the nonlinear least squares problem (2.9) with *I t*_*V,j*_; *V*_*C,i*_; *p* given by the function (2.28) with *V*_*C*_ = *V* _*C,i*_ at times *t*_*V,j*_ *∈* [*t*_*V*,0_, *t*_*V,f*_ ), and *I*_meas_ (*t*_*V,j*_ ; *V*_*C,i*_) the measured currents.

#### Assumption 2.14.

*Assume that n*_*V*_ *>* 1, *the sample times* 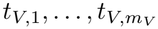 *are distinct with m*_*V*_ *≥* 15, *V*_*C,i*_ *≠V*_*C,j*_ *for i ≠ j, and the parameters* (2.29) *satisfy* 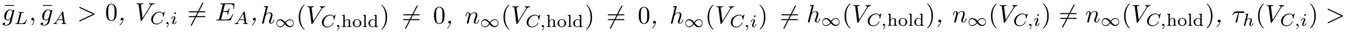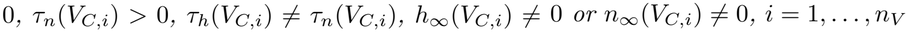.

The following theorem shows that the rank of the associated Jacobian is equal to *n*_*P*_ *−* 2. For simplicity, we omit the case *n*_*V*_ = 1.

#### Theorem 2.15.

*Let* 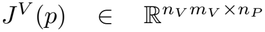 *denote the Jacobian corresponding to* (2.9) *with* (*I t*_*V,j*_; *V*_*C,i*_; *p*) *given by the function* (2.28) *and with V*_*C*_ = *V*_*C,i*_ *at times t*_*V,j*_ *∈* [*t*_*V*,0_, *t*_*V,f*_ ). *If Assumption 2*.*14 holds then J*^*V*^ (*p*) *has rank n*_*P*_ *−* 2 = 4 + 4*n*_*V*_ .

*Proof* The proof is given in Appendix A.6. Actually, the statement of this theorem holds with power 3 in the current equation (2.28) replaced by any positive integer *q ∈* ℕ. In Appendix A.6, we will prove the more general result for power 3 replaced by *q ∈* ℕ.

The proof of Theorem 2.15 shows that the vectors in the null space of *J*^*V*^ (*p*) correspond to the parameter degeneracy due to the multiplicative combination of 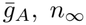 and *h*_*∞*_ in (2.28). Fixing *h*_*∞*_(*V*_*C*,hold_) = 1 and 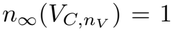 so that *ñ*_*P*_ = *n*_*P*_ *−* 2 results in a full rank Jacobian.

#### Corollary 2.16.

*Let Assumption 2*.*14 hold. The Jacobian* 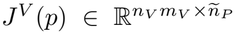*corresponding to* (2.9) *with parameters* (2.29) *and* 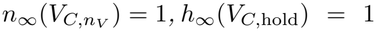 *has full column rank, ñ* _*P*_ =4 + 4*n*_*V*_ .

*Proof* The proof follows from Theorem 2.15 and the fact that the columns of the Jacobian of (2.9) corresponding to the partial derivative with respect to 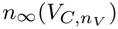 and *h*_*∞*_(*V*_*C*,hold_) can be expressed as linear combinations of the other columns, as is shown in the proof of Theorem 2.15 in Appendix A.6.

These results are illustrated numerically in Section 3 (Fig. 4B).

### 2.5 Action Potential Generating Currents

The action potential is generated by two ionic currents, a fast-activating and inactivating sodium current whose opening leads to the subsequent delayed activation of a potassium current, called the delayed rectifier current. The delayed rectifier current contributes to the repolarization of the membrane potential toward its resting value. In the Numerics Section 3, we consider the fast sodium and delayed rectifier currents modeled from STG LP neuron recordings (Liu et al. 1998), see Fig. 5A, C.

**Fig. 5.**
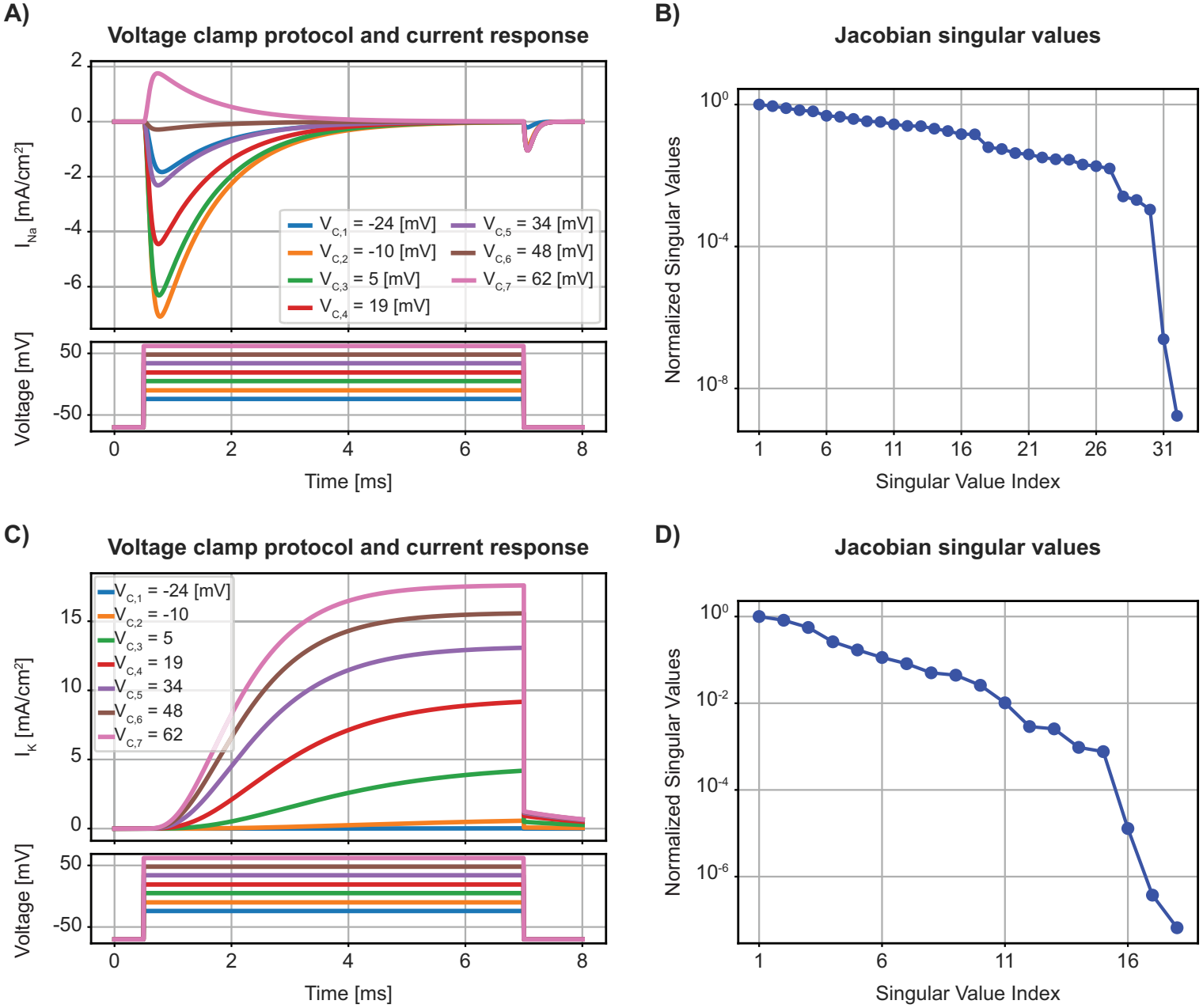
Parameter degeneracy in voltage clamp experiments for STG sodium and potassium current models: **(A)** Sodium voltage clamp traces for the sodium current model. **(B)** Normalized singular values of the Jacobian evaluated at the corresponding model parameters. **(C)** Potassium voltage clamp traces for the potassium current model. **(D)** Normalized singular values of the Jacobian evaluated at the corresponding model parameters.

#### 2.5.1 Sodium current

The model follows the form presented in (2.1) – (2.7) with Γ = *{L, Na}, p*_*L*_ = *q*_*L*_ = 0, *p*_*Na*_ = 3, *q*_*Na*_ = 1, *a*_*Na*_ = *m*, and *b*_*Na*_ = *h*.

Using the analytical solutions for the gating variables as in (2.10), the equation for the current is

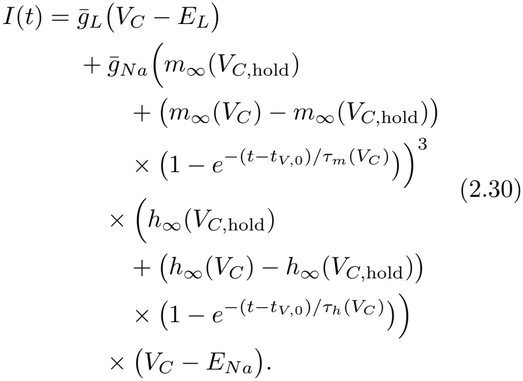

Again, we have *n*_*V*_ voltage clamp recordings with command voltages *V*_*C,i*_, *i* = 1, …, *n*_*V*_, and estimate the unknown parameters

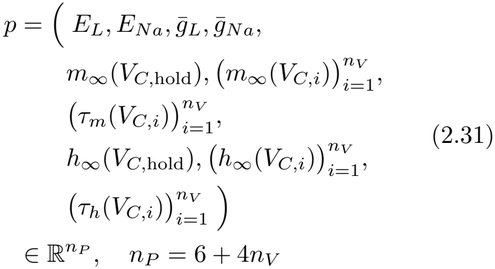

as the solution of the nonlinear least squares problem (2.9) with *I* (*t*_*V,j*_; *V*_*C,i*_; *p*) given by the function (2.30) with 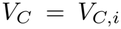 at times *t*_*V,j*_ *∈* [*t*_*V*,0_, *t*_*V,f*_ ), and *I*_meas_ (*t*_*V,j*_ ; *V*_*C,i*_) the measured currents. Because the functional form of (2.30) is identical to (2.28), Theorem 2.15 applies and the associated Jacobian has rank *n*_*P*_ *−* 2 = 4 + 4*n*_*V*_ .

The model-based multiplicative parameter degeneracy in (2.30) is usually resolved by fixing the value of *h*_*∞*_(*V*_*C*,hold_) (e.g., = 1 as in Liu et al. 1998), and by assuming that 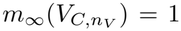 at the largest depolarizing command voltage, 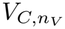. Again, Corollary 2.16 applies and the associated Jacobian is full rank. This result is illustrated numerically in Section 3 (Fig. 5B, blue line and data points).

#### 2.5.2 Potassium current

The model follows the form presented in (2.1) – (2.7) with Γ = *{L, K}, p*_*L*_ = *q*_*L*_ = 0, *p*_*K*_ = 4, *q*_*K*_ = 0, and *a*_*K*_ = *n*.

Using the analytical solutions for the gating variables as in (2.10), the equation for the current is

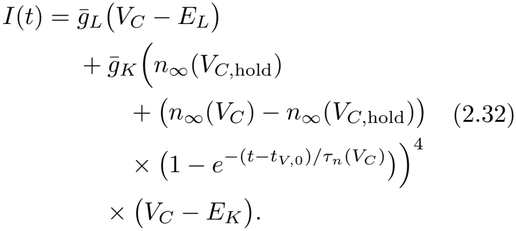

The voltage clamp protocol is identical to that of the sodium voltage clamp experiment, see Fig. 5C. For *n*_*V*_ voltage clamp recordings with command voltages *V*_*C,i*_, *i* = 1, …, *n*_*V*_, we estimate

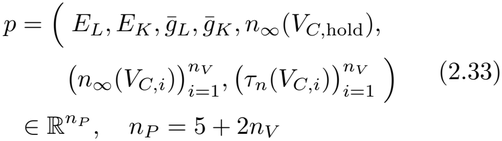

as the solution of the nonlinear least squares problem (2.9) with *I* (*t*_*V,j*_; *V*_*C,i*_; *p*) given by the function (2.32) with *V*_*C*_ = *V*_*C,i*_ at times *t*_*V,j*_ *∈* [*t*_*V*,0_, *t*_*V,f*_ ), and *I*_meas_ (*t*_*V,j*_; *V*_*C,i*_) the measured currents.

##### Assumption 2.17.

*Assume that n*_*V*_ *>* 1, *the sample times* 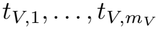 *are distinct with m*_*V*_ *≥* 9, *V*_*C,i*_ ≠ *V*_*C,j*_ *for i≠j, and the parameters* (2.33) *satisfy* 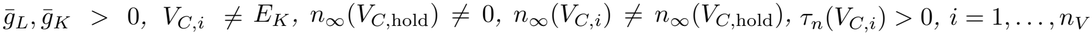.

The following theorem shows that the rank of the associated Jacobian is equal to *n*_*P*_ *−* 1. For simplicity, we omit the case *n*_*V*_ = 1.

##### Theorem 2.18.

*Let* 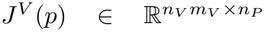 *denote the Jacobian corresponding to* (2.9) *with I* (*t*_*V,j*_; *V*_*C,i*_; *p*) *given by the function* (2.32) *and with V*_*C*_ = *V*_*C,i*_ *at times t*_*V,j*_ *∈* [*t*_*V*,0_, *t*_*V,f*_ ). *If Assumption 2*.*17 holds then J*^*V*^ (*p*) *has rank n*_*P*_ *−* 1 = 4 + 2*n*_*V*_ .

*Proof* The proof is given in Appendix A.7. Actually, the statement of this theorem holds with power 4 in the current equation (2.32) replaced by any integer *q ≥* 2. In Appendix A.7, we will prove the more general result for power 4 replaced by *q ≥* 2.

Proceeding as above to eliminate the multiplicative parameter degeneracy in (2.32) by setting 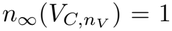 at the largest voltage clamp step, 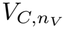, so that *ñ*_*P*_ = *n*_*P*_ *−*1, leads to the following corollary.

##### Corollary 2.19.

*Let Assumption 2*.*17 hold. The Jacobian* 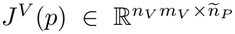*corresponding to* (2.9) *with parameters* (2.33) *and* 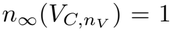*has full column rank ñ*_*P*_ = 4 + 2*n*_*V*_ .

*Proof* The proof follows from Theorem 2.18 and the fact that the column of the Jacobian of (2.9) corresponding to the partial derivative with respect to 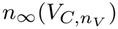 can be expressed as a linear combination of the other columns, as is shown in the proof of Theorem 2.18 in Appendix A.7.

The result in Corollary 2.19 is illustrated numerically in Section 3 (Fig. 5D, blue line and data points).

## 3 Numerics

We provide numerical examples of the theoretical results derived in Section 2 using ion current models derived from STG LP neuron recordings, and from cerebellar Purkinje cell recordings (Prinz et al. 2003; Liu et al. 1998; Angelo et al. 2007; Golowasch and Marder 1992).

### 3.1 Whole Cell H-Current Model

A typical voltage clamp experiment consists of five voltage clamp steps below the resting potential, see Fig. 2A. Based on these data, Corollary 2.3*i*. predicts a Jacobian rank deficiency of two, which is verified numerically in Fig. 2B (blue data points and connecting line). Next, Corollary 2.6*i*. predicts that addition of the tail current experiment, see Fig. 2C, reduces the rank deficiency by one, as confirmed in Fig. 2B (red data points and connecting line). Finally, Corollary 2.9 states that addition of a current clamp experiment after block of the H-current, see Fig. 2D, results in a full rank Jacobian. This is indeed the case in our numerical example, see Fig. 2B (green data points and connecting line).

### 3.2 Cell-Attached H-Current Model

A typical voltage clamp experiment on cerebellar Purkinje cells in the cell-attached configuration consists of seven voltage clamp steps below the resting potential, see Fig. 3A. In this example, the activation function *h*_*∞*_ at the holding potential is equal to zero, see Fig. 3B (red dashed line and dot). Based on these data, Corollary 2.11*i*.*a*. predicts a Jacobian rank deficiency of one, which is verified numerically in Fig. 3C (red data points and connecting line). In contrast, if the activation function is not equal to zero at *V*_*C*,hold_, Corollary 2.11*i*.*b*. predicts a full rank Jacobian. We thus consider a slight modification of the activation function as in Fig. 3B (blue line and dot). In this case, the Jacobian indeed has full column rank, see Fig. 3C (blue data points and connecting line). Finally, Corollary 2.13 predicts that adding the tail portion of the voltage clamp recording, see Fig. 3D, results in a full rank Jacobian even if *h*_*∞*_(*V*_*C*,hold_) = 0. This is confirmed in Fig. 3C (green data points and connecting line).

### 3.3 A-Type Potassium Current Model

A typical voltage clamp experiment consists of five depolarizing voltage clamp steps from a hyperpolarized holding potential, see Fig. 4A. Based on these data, Theorem 2.15 predicts a Jacobian rank deficiency of two, which is verified numerically in Fig. 4B (blue data points and connecting line). Next, fixing *h*_*∞*_(*V*_*C*,hold_) = 1 reduces the rank deficiency by one, as confirmed in Fig. 4B (green data points and connecting line). Finally, Corollary 2.16 states that additionally fixing 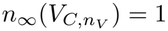 results in a full rank Jacobian. This is indeed the case in our numerical example, see Fig. 4B (red data points and connecting line).

### 3.4 Action Potential Generating Current Models

A typical voltage clamp experiment for the sodium current consists of seven depolarizing voltage clamp steps; see Fig. 5A. Since the sodium current model has the same functional form as the A-current model, Theorem 2.15 predicts a Jacobian rank deficiency of two. After fixing *h*_*∞*_(*V*_*C*,hold_) = 1 and 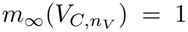, Corollary 2.16 predicts a full rank Jacobian, which is verified numerically in Fig. 5B (blue data points and connecting line). The potassium current is recorded using the same voltage clamp protocol; see Fig. 5C. In this case, Theorem 2.18 predicts a Jacobian rank deficiency of one, and Corollary 2.19 states that fixing 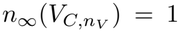 results in a full rank Jacobian. This is indeed the case in our numerical example; see Fig. 5D (blue data points and connecting line).

## 4 Discussion

Jacobians play a central role in the solution of nonlinear least squares problems (see, e.g., Björck 2024, Chap. 8; Dennis and Schnabel 1996, Chap. 10; Nocedal and Wright 2006, Chap. 10), and in proving the existence of local solution manifolds and their computation (see, e.g., Fracasso and Heinkenschloss 2027; Eriksson et al. 2005; Namazifard et al. 2026). In this work, we analyzed the structure of the Jacobians associated with a classical problem in biophysics, the fitting of HH-type current models to voltage clamp data (Hille 2001, Chap. 2). Our results show that the column vectors of these Jacobians are never linearly independent. One fundamental assumption of HH-type models is that the peak conductance multiplies the activation and inactivation variables, if present. This inevitably leads to linear dependence between the Jacobian column vectors and thus to a local manifold of equivalent solutions.

Even after fixing this simple form of parameter degeneracy, a local manifold of equivalent solutions may still exist. Using a combination of numerical simulations and analytical work, we established that the simplest HH-type model with one activation variable, such as that used to describe H-currents, has a non-trivial solution manifold when only voltage clamp data is available. This result is independent of the model’s specific parameter values. We also show how additional experiments can eliminate this parameter degeneracy. It is worth pointing out that voltage clamp experiments are notoriously difficult and hence these additional experiments might not always be technically feasible. The experimental setting of cell-attached patch clamp recordings also reveals an unexpected parameter degeneracy when the channel is closed at the holding potential. Note that this condition is often fulfilled in published cell-attached recordings of the H-current (Magee 1998; Angelo et al. 2007).

Our results show that more complex HH-type models do not exhibit parameter degeneracy beyond that caused by the model formulation discussed above. However, the singular value spectrum of their Jacobians often spans more than six orders of magnitude, indicative of ill-conditioning (Hansen 1998). The implications of this result for parameter estimation are analyzed in a companion paper (Namazifard et al. 2026). In the same paper, another interesting issue is addressed: how to trace numerically a solution manifold away from a given parameter set when its existence is established.

In contrast to the simple example discussed in the introduction, the results presented there show that HH-type model solution manifolds are nonlinear, involving multiple co-varying parameters. Importantly, we expect that the results presented in these two papers will generalize to situations where the membrane potential of a compartmental model is not clamped to a fixed value (Namazifard et al. 2026).

We are not aware of any application of Jacobians to study model parameter degeneracy. Similarly, simultaneous parameter estimation has been considered in the past (Willms et al. 1999), but mainly from a numerical perspective; the same holds for model-based multiplicative parameter degeneracy (Csercsik et al. 2012; Walch and Eisenberg 2016). Our results go well beyond these earlier considerations, by fully characterizing the rank structure and the null space of HH-type current model Jacobians under the relevant range of experimental conditions encountered in neurophysiological studies.

## Declarations

### Data availability

Code for reproducing all figures and synthetic data in this paper is available at [https://doi.org/10.5281/zenodo.24204791]. The DOI will become publicly accessible upon publication of the manuscript.

### AI use

Parts of the proofs in Appendices A.6 and A.7 were initially obtained with the help of Claude (Anthropic), and then reviewed and edited by the authors. The authors take full responsibility for the content of the published article.

### Author contributions

All authors participated in the conceptualization of the work; S.N. carried out the numerical work; M.H. implemented hand proofs; F.G. implemented AI for parts of proofs using Claude (Anthropic); M.H. and F.G. wrote and edited the manuscript.

## Appendix A Proofs of the Rank Results for Residual Jacobians

### A.1 Proof of Theorem 2.2 (rank *J*^*V*^ (*p*); H-current, whole cell)

This section provides a proof of the theorem on the Jacobian rank of the H-current model constrained by the voltage clamp experiment. We introduce the abbreviations

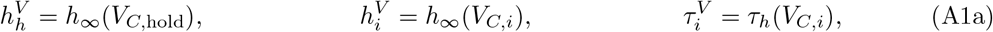

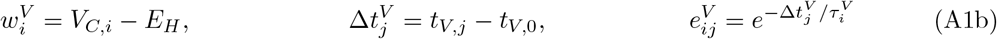

for *j* = 1, …, *m*_*V*_, *i* = 1, …, *n*_*V*_ . Furthermore, for each experiment *i* = 1, …, *n*_*V*_, we define the 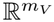 vectors **1** = (1, …, 1)^*T*^, **0** = (0, …, 0)^*T*^,

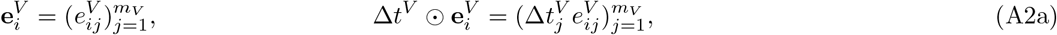

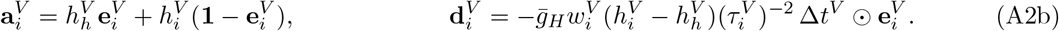

With this notation, the current (2.10) at *V*_*C*_ = *V*_*C,i*_ and *t* = *t*_*V,j*_ is written as

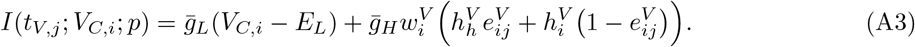

The residual 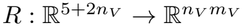 is a vector of *n*_*V*_ blocks, each of length *m*_*V*_, and has components

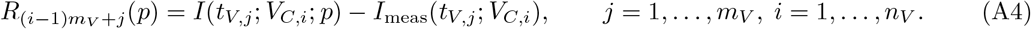

The partial derivatives of (A4), for component *j* of block *i* and *k* = 1, …, *n*_*V*_ are

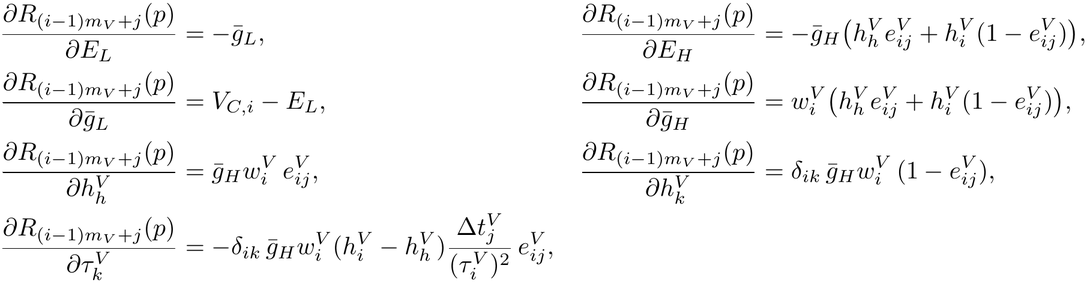

where *δ*_*ik*_ is the Kronecker delta, i.e., *δ*_*ik*_ = 0 if *i ≠ k* and *δ*_*ii*_ = 1, and where we used 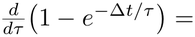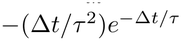. The Jacobian in block form is

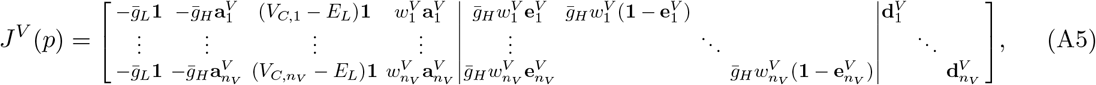

with the *i*-th row block corresponding to voltage clamp *V*_*C,i*_, and with columns ordered as in (2.11):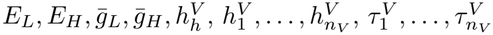.

i. Let *n*_*V*_ *>* 1. Using the expression (A5) of the Jacobian one can check that

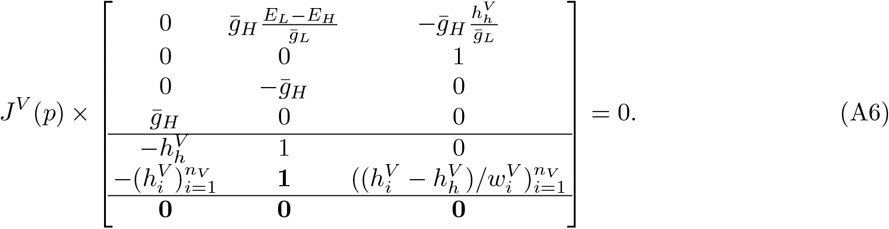 The three column vectors in (A6) are linearly independent. Thus, the rank of the Jacobian *J*^*V*^ (*p*) is at most 2 + 2*n*_*V*_ . Note that the first column vector in (A6), which written in full notation is 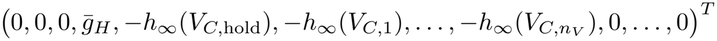, reflects the fact that 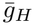 appears only in a product with 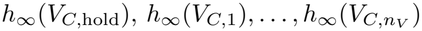 in (2.10) and, therefore, these parameters can never be identified simultaneously. Next, we will show that the 2 + 2*n*_*V*_ columns 1, 3, 6, …, 5 + 2*n*_*V*_ of the Jacobian (A5) are linearly independent, which implies that the Jacobian (A5) has rank 2 +2*n*_*V*_ . We set a linear combination of these columns to zero and show that the coefficients 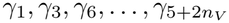 in this linear combination must be zero. Consider

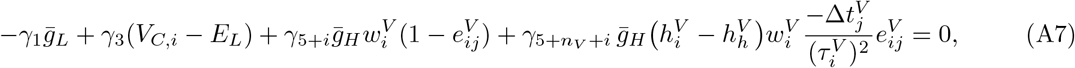

for *j* = 1, …, *m*_*V*_ and *i* = 1, …, *n*_*V*_ . For fixed experiment index *i*, subtracting the *j* = 1 equation in (A7) from the others, then dividing by 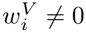 gives

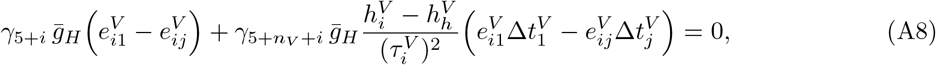

*i* = 1, …, *n*_*V*_, *j* = 2, …, *m*_*V*_ . Because for *t*_*V,j*_ *≠ t*_*V,ℓ*_, the 2 *×* 2 matrix

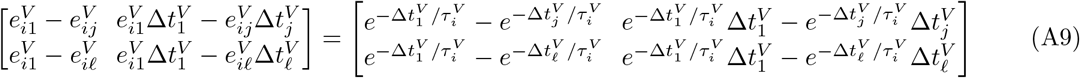

is nonsingular, (A8) and 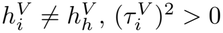 imply *γ*_5+*i*_ = *γ*_5+*nv*+*i*_ = 0, *i* = 1, …, *n*_*V*_ . Hence

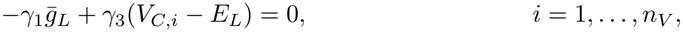

which, because of *n*_*V*_ *>* 1 and *V*_*C,i*_ *≠ V*_*C,j*_ for *i ≠ j*, implies *γ*_1_ = *γ*_3_ = 0.
ii. Let *n*_*V*_ = 1. The parameters are 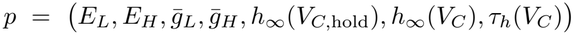 . Using the expression (A5) of the Jacobian, one can check that

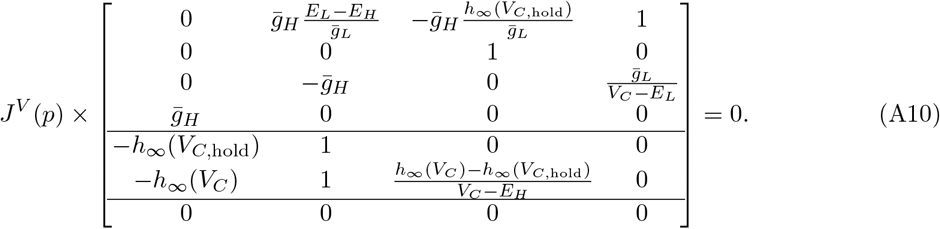

Thus, the rank of the Jacobian is at most 3. Proceeding as in part i, one can show that columns 1, 6 and 7 of (A5) are linearly independent, which implies that the rank of the Jacobian is 3.

#### A.2 Proof of Theorem 2.5 (rank *J*^*V T*^ (*p*); H-current, whole cell)

This section provides a proof of the theorem on the Jacobian rank of the H-current model constrained by the voltage clamp and tail current experiment. Analogously to (A1) and (A2), we introduce the abbreviations

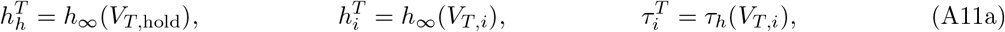

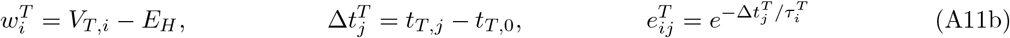

for *j* = 1, …, *m*_*T*_, *i* = 1, …, *n*_*T*_ . Furthermore, for each experiment *i* = 1, …, *n*_*T*_, we define the 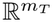 vectors

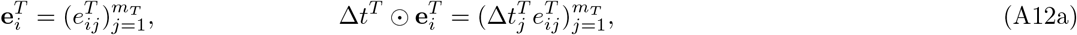

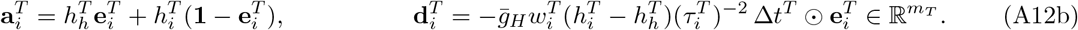

Rather than introducing two different notations, we let **1** = (1, …, 1)^*T*^ and **0** = (0, …, 0)^*T*^ denote vectors in 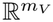 or in 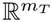, with either size depending on the context in which they are used.

We write the Jacobian (A5) as

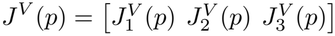

with 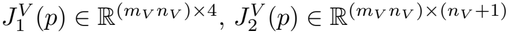, and 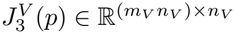.

Adding tail current recordings results in additional parameters in (2.15) and additional components in the residual function corresponding to the second double sum in (2.16). The Jacobian of the residual function corresponding to (2.16) is given by

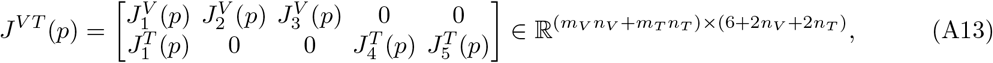

where the partial Jacobians 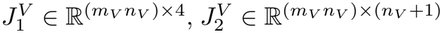, and 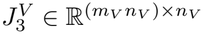 correspond to the voltage clamp recordings. The matrices in the second row block correspond to tail current recordings. Specifically, 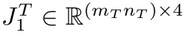 is the partial Jacobian of derivatives with respect to 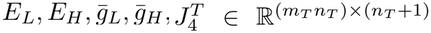 is the partial Jacobian of derivatives with respect to 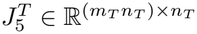 is the partial Jacobian of derivatives with respect to 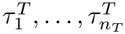. The partial Jacobians corresponding to the tail current recordings are

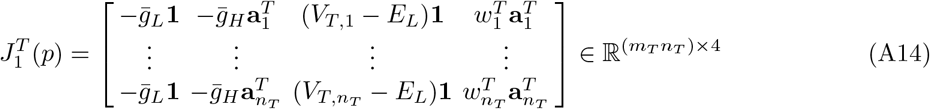

which is the partial Jacobian of derivatives with respect to 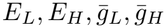,

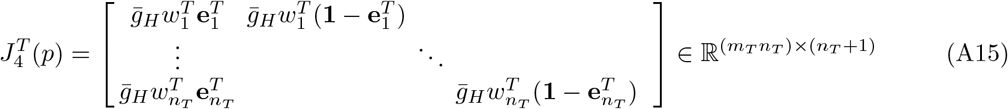

which is the partial Jacobian of derivatives with respect to 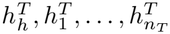, and

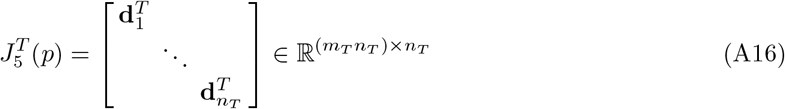

which is the partial Jacobian of derivatives with respect to 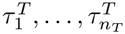.

i. Let *n*_*T*_ *>* 1. From the proof of Theorem 2.2 in Appendix A.1 we know that 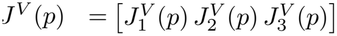 and 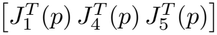 each have null spaces of dimension 3 and that (A6) and

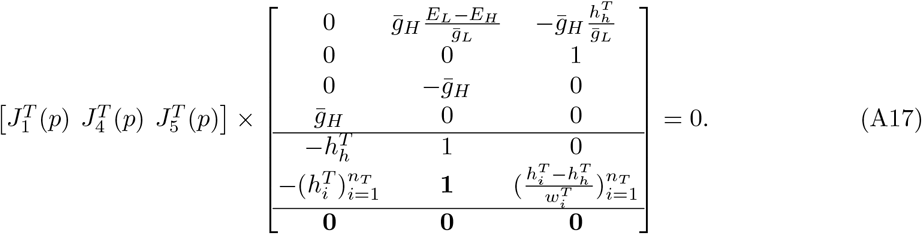 Because of the block structure of the Jacobian (A13) and (A6), (A17), a vector 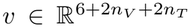 is in the null space of the Jacobian (A13) if and only if there exist 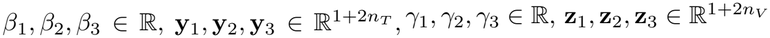 such that

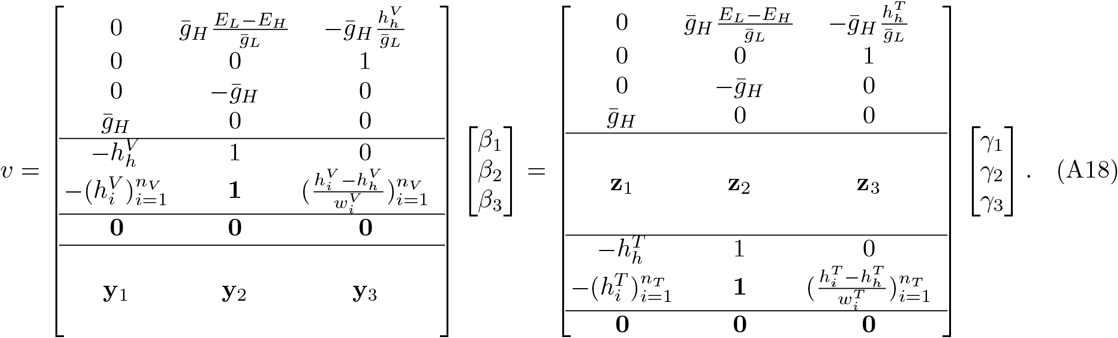 Because of the first four rows in (A18) and 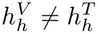, a vector 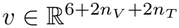 satisfies (A18), i.e., is in the null space of the Jacobian (A13), if and only if it is in the span of the two vectors

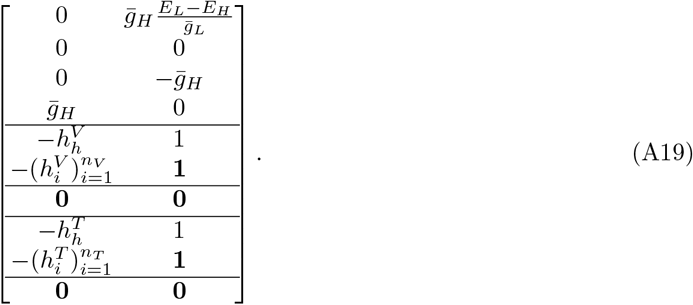 The first column vector in (A19) corresponds to *β*_1_ = 1, *β*_2_ = *β*_3_ = 0, *γ*_1_ = 1, *γ*_2_ = *γ*_3_ = 0 in (A18), and the **y**’s and **z**’s chosen accordingly; the second column vector in (A19) corresponds to *β*_2_ = 1, *β*_1_ = *β*_3_ = 0, *γ*_2_ = 1, *γ*_1_ = *γ*_3_ = 0, and the **y**’s and **z**’s chosen accordingly. Because the column vectors in (A19) are a basis for the null space of the Jacobian (A13), the rank of the Jacobian (A13) is 4 + 2*n*_*V*_ + 2*n*_*T*_ . Note that the first column vector in (A19) reflects the fact that 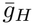 appears only in a product with 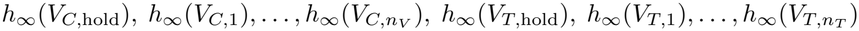 in (2.10) and, therefore, these parameters can never be identified simultaneously. If 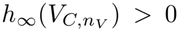, the (5 + *n*_*V*_ )-th column corresponding to the partial derivative with respect to 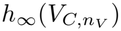 in the Jacobian (A13) can be expressed using the other columns.
ii. Let *n*_*T*_ = 1. From the proof of Theorem 2.2 in Appendix A.1 we know that 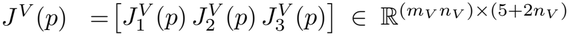 has a null space of dimension 3 and (A6) holds, and that 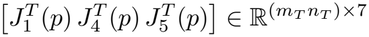 has a null space of dimension 4 and

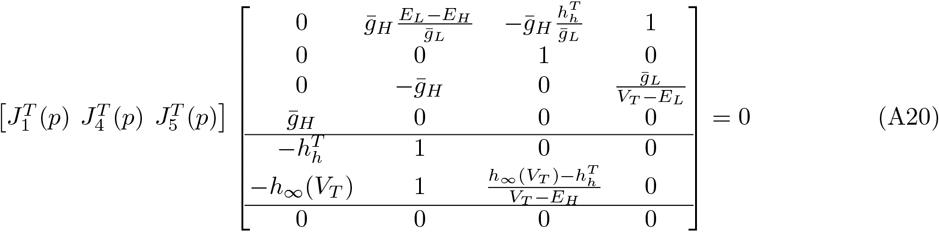

(see (A10)). Because of the block structure of the Jacobian (A13) and (A6), (A20), a vector 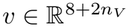 is in the null space of the Jacobian (A13) if and only if there exist 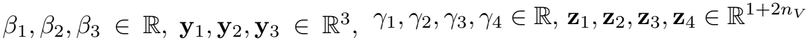 such that

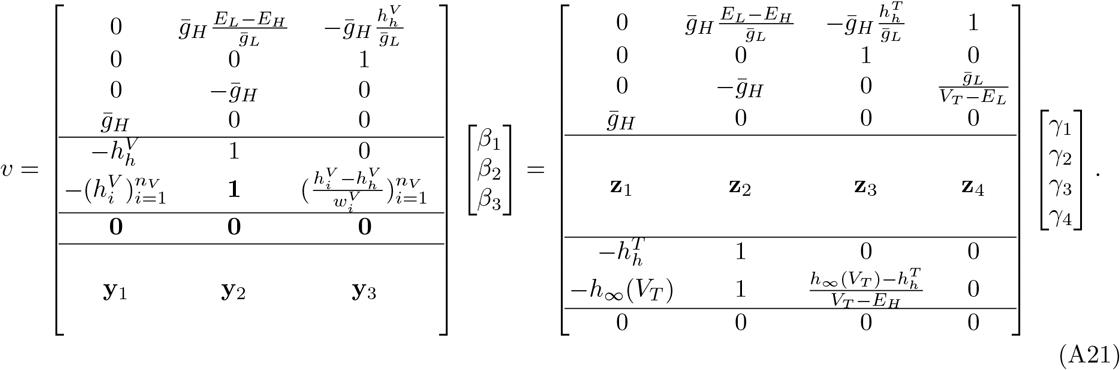 Because of the first four rows in (A21) and 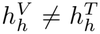, a vector 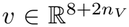 satisfies (A21), i.e., is in the null space of the Jacobian (A13), if and only if it is in the span of the three vectors

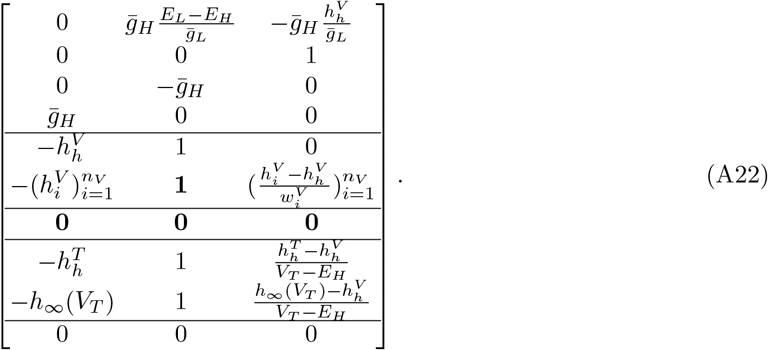

The first column vector in (A22) corresponds to *β*_1_ = 1, *β*_2_ = *β*_3_ = 0, *γ*_1_ = 1, *γ*_2_ = *γ*_3_ = *γ*_4_ = 0 in (A21), and the **y**’s and **z**’s chosen accordingly; the second column vector in (A22) corresponds to *β*_2_ = 1, *β*_1_ = *β*_3_ = 0, *γ*_2_ = 1, *γ*_1_ = *γ*_3_ = *γ*_4_ = 0, and the **y**’s and **z**’s chosen accordingly; the third column vector in (A22) corresponds to *β*_1_ = *β*_2_ = 0, *β*_3_ = 1,

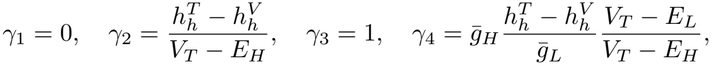

and the **y**’s and **z**’s chosen accordingly. The column vectors in (A22) are linearly independent and are therefore a basis for the null space of the Jacobian (A13). Hence, the rank of the Jacobian (A13) is 5 + 2*n*_*V*_ . Again, note that the first column vector in (A22) reflects the fact that 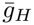 appears only in a product with 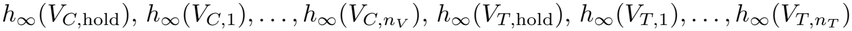 in (2.10) and, therefore, these parameters can never be identified simultaneously. If 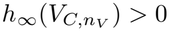, the (5 + *n*_*V*_ )-th column corresponding to the partial derivative with respect to 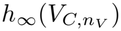 in the Jacobian (A13) can be expressed using the other columns.

#### A.3 Proof of Theorem 2.8 (rank *J*^*V T C*^(*p*); H-current, whole cell)

This section provides a proof of the theorem on the Jacobian rank of the H-current model constrained by the voltage clamp, tail current and current clamp experiment. Recall that the parameters are given by (2.15). The Jacobian of the residual function corresponding to (2.20) is given by

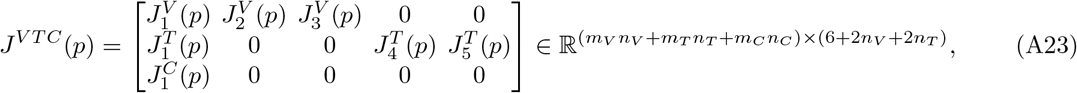

where the partial Jacobians 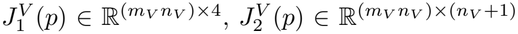, and 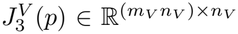 correspond to the voltage clamp recordings and the partial Jacobians 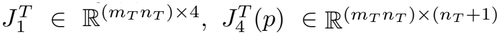, and 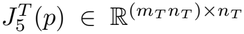 correspond to the tail current recordings. These partial Jacobians are the ones in (A13).

The partial Jacobian 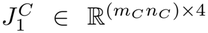 is the partial Jacobian of derivatives with respect to 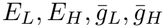 corresponding to the current clamp recordings. Defining the vectors

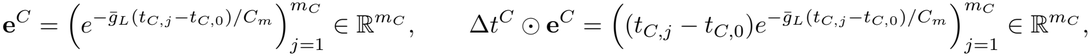

this partial Jacobian is given by

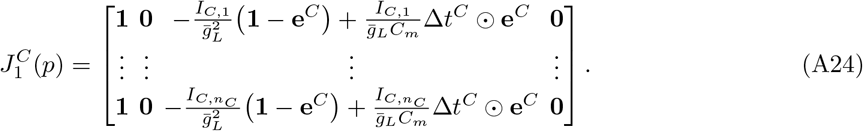

i. Let *n*_*T*_ *>* 1. In Section A.2 it was shown that the two columns of (A19) are a basis for the null space of (A13), the submatrix of (A23) consisting of the first *m*_*V*_ *n*_*V*_ + *m*_*T*_ *n*_*T*_ rows. The first column vector in (A19) is also in the null space of

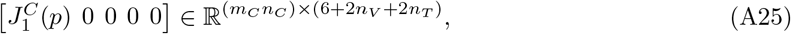

while the second column vector in (A19) is not. Hence, (A23) has rank 5 + 2*n*_*V*_ + 2*n*_*T*_ . As already mentioned in Section A.2, the first column vector in (A19) reflects the fact that 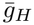 appears only in a product with 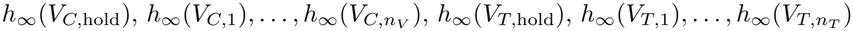 in (2.10) and, therefore, these parameters can never be identified simultaneously. If 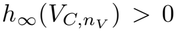, the (5 + *n*_*V*_ )-th column corresponding to the partial derivative with respect to 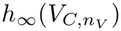 in the Jacobian (A23) can be expressed using the other columns.
ii. Let *n*_*T*_ = 1. In Section A.2 it was shown that the three columns of (A22) are a basis for the null space of (A13), the submatrix of (A23) consisting of the first *m*_*V*_ *n*_*V*_ + *m*_*T*_ *n*_*T*_ rows. The first column vector in (A22) is also in the null space of (A25), while the second and third column vectors in (A22) are not. Hence, (A23) has rank 7 + 2*n*_*V*_ . As already mentioned in Section A.2, the first column vector in (A22) reflects the fact that 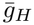 appears only in a product with 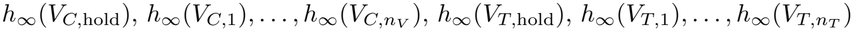 in (2.10) and, therefore, these parameters can never be identified simultaneously.

#### A.4 Proof of Theorem 2.10 (rank *J*^*V*^ (*p*); H-current, cell-attached)

This section provides a proof of the theorem on the Jacobian rank of the isolated H-current model constrained by the voltage clamp experiment. We use the abbreviations (A1) and (A2) introduced in Appendix A.1. As in Appendix A.1, the Jacobian 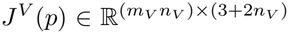 of the residual function is given by

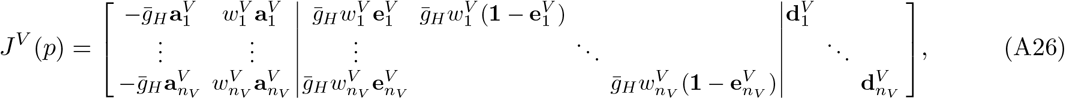

with the *i*-th row block corresponding to voltage clamp *V*_*C,i*_, and with columns ordered as in (2.22):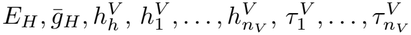.

i. Let *n*_*V*_ *>* 1. We consider the two cases 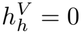 and 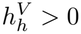.
i.a. The first case is 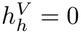. In this case, one can verify that

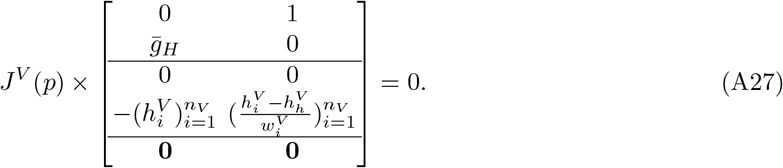 The null space vectors in (A27) are linearly independent. Hence, if 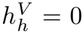, the rank of the Jacobian is at most 1 + 2*n*_*V*_ . Next, we show that the columns 3, …, 3 + 2*n*_*V*_ of the Jacobian (A26) are linearly independent, which implies that the Jacobian (A26) has rank 1 + 2*n*_*V*_ . Setting the linear combination of columns 3, …, 3 + 2*n*_*V*_ of the Jacobian (A26) to zero gives

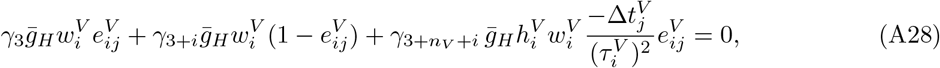

for *j* = 1, …, *m*_*V*_ and *i* = 1, …, *n*_*V*_ . We will show that 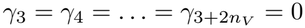. For fixed experiment index *i*, subtracting the *j* = 1 equation in (A28) from the others and then dividing by 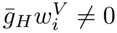 gives

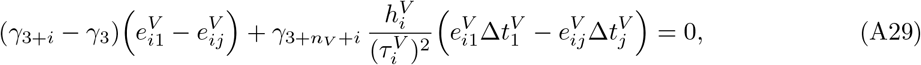

*i* = 1, …, *n*_*V*_, *j* = 2, …, *m*_*V*_ . Because for *t*_*V,j*_ *≠ t*_*V,ℓ*_, the 2 *×* 2 matrix (A9) is nonsingular, (A29) and the assumption 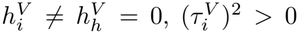 imply 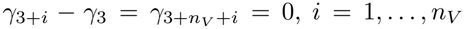. Inserting these identities into (A28) gives 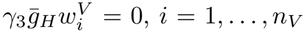, which implies *γ*_3_ = 0. Hence 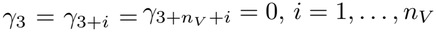.
i. b. The second case is 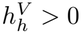. In this case, one can verify that

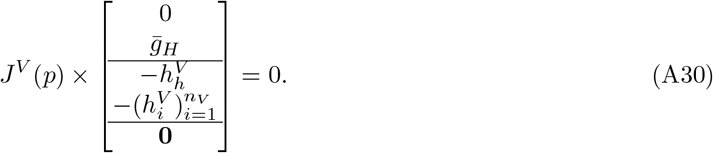

Hence, if 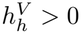, the rank of the Jacobian is at most 2 + 2*n*_*V*_.

Next, we show that the columns 1, 3, …, 3 + 2*n*_*V*_ of the Jacobian (A26) are linearly independent, which implies that the Jacobian (A26) has rank 2 + 2*n*_*V*_ . Again, we set a linear combination of these columns to zero and show that the coefficients 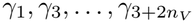 in this linear combination must be zero. Setting the linear combination of columns 1, 3, …, 3 + 2*n*_*V*_ of the Jacobian (A26) to zero gives

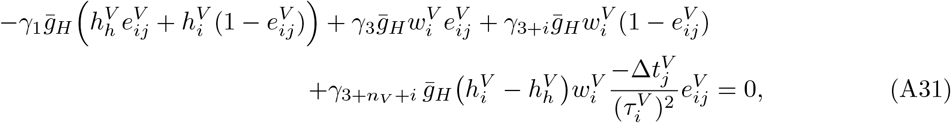

for *j* = 1, …, *m*_*V*_ and *i* = 1, …, *n*_*V*_ .

For fixed experiment index *i*, subtracting the *j* = 1 equation in (A31) from the others and then dividing by 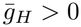 gives

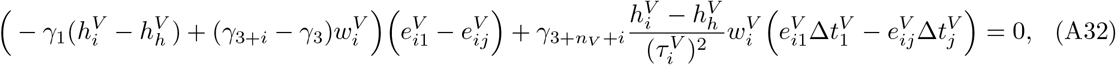

*i* = 1, …, *n*_*V*_, *j* = 2, …, *m*_*V*_ . Because for *t*_*V,j*_ *≠ t*_*V,ℓ*_, the 2 *×* 2 matrix (A9) is nonsingular, (A32) and the assumption 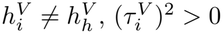 imply

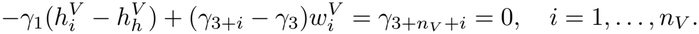

Inserting these identities into (A31) and dividing by 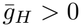 gives

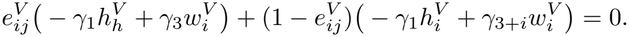

By the first identity above, the two expressions in parentheses are equal, and therefore both vanish:

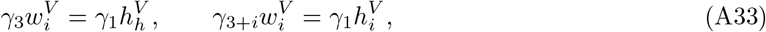

for *i* = 1, …, *n*_*V*_ . Because *n*_*V*_ *>* 1 and the command voltages are distinct, 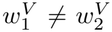, and subtracting the first equation in (A33) for *i* = 2 from the one for *i* = 1 gives 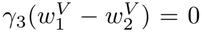. Then 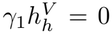 and 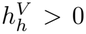 imply *γ*_1_ = 0, and the second equation in (A33) with 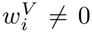 gives *γ*_3+*i*_ = 0. Altogether, 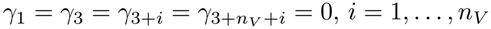.

ii. Let *n*_*V*_ = 1. The Jacobian is

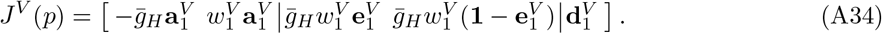 One can check that

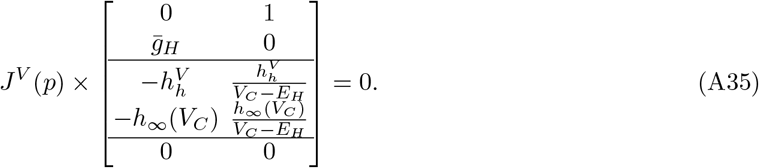

Thus, the rank of the Jacobian (A34) is at most 3.

As in part i.a. of the proof, one can show that columns 3, 4, 5 of (A34) are linearly independent, which implies that the rank of the Jacobian is 3.

#### A.5 Proof of Theorem 2.12 (rank *J*^*V T*^ (*p*); H-current, cell-attached)

This section provides a proof of the theorem on the Jacobian rank of the isolated H-current model constrained by the voltage clamp and tail current experiment. We use the abbreviations (A1) and (A2) introduced in Appendix A.1. In addition define the abbreviations

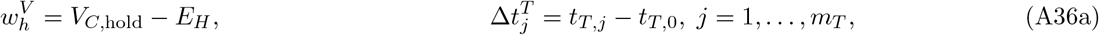

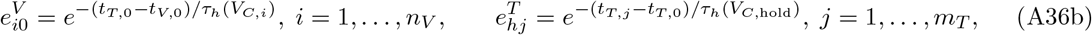

and the vectors

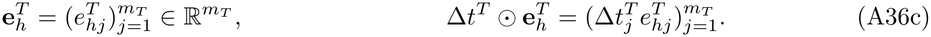

We write the Jacobian in (A26) as 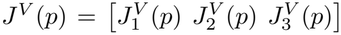 with partial Jacobians 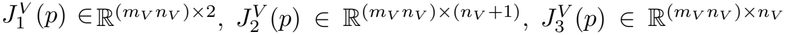. The Jacobian of the residual function corresponding to (2.24) is given by

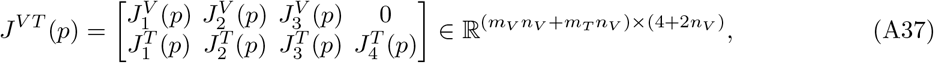

where 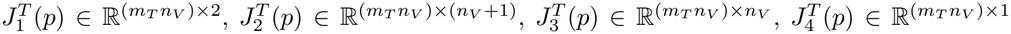, are the partial Jacobians of the residual terms in the second double sum in (2.24) with respect to 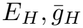, to 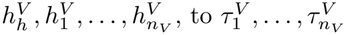, and to *τ*_*h*_(*V*_*C*,hold_), respectively. We will derive them next.

Using (A1) and (A36), (2.25), (2.26) are written as

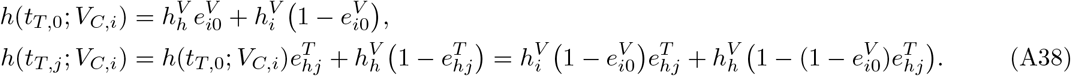

Using (A38) in (2.27) gives

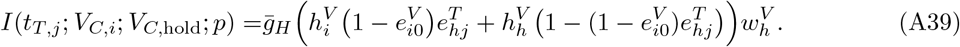

From this last expression the partial Jacobians corresponding to the tail current experiment are

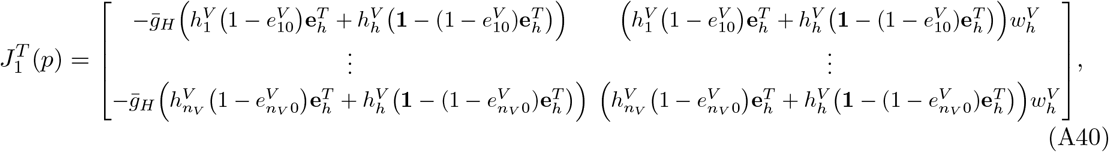

and

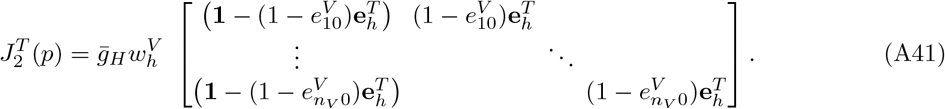

Moreover, using (A39) and 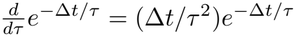 one obtains

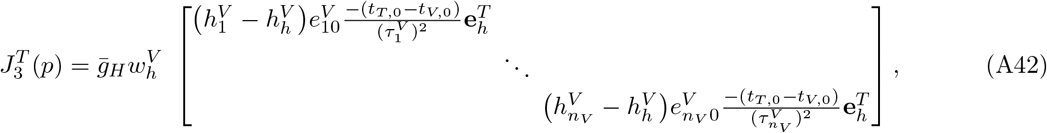

and

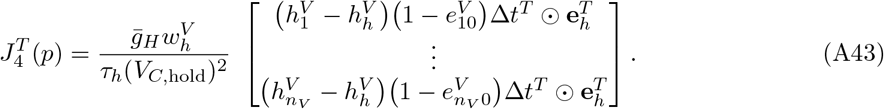

The null space of the Jacobian *J*^*V T*^ (*p*) in (A37) is

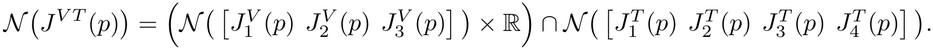

i.a. Let *n*_*V*_ *>* 1 and 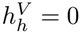. By (A27),

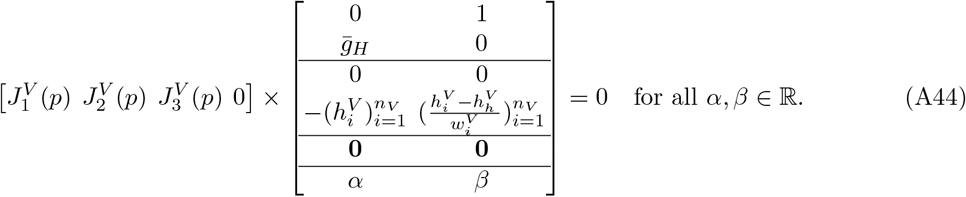

From (A40)–(A43) one can check that the first column vector in (A44) is in 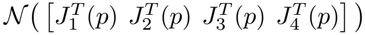 if and only if *α* = 0, and that the second column vector in (A44) is not in 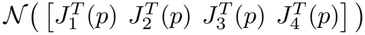 for any *β ∈* ℝ . Hence, the rank of Jacobian *J*^*V T*^ (*p*) in (A37) is 3 + 2*n*_*V*_ .

i. b. Let *n*_*V*_ *>* 1 and 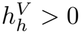. By (A30),

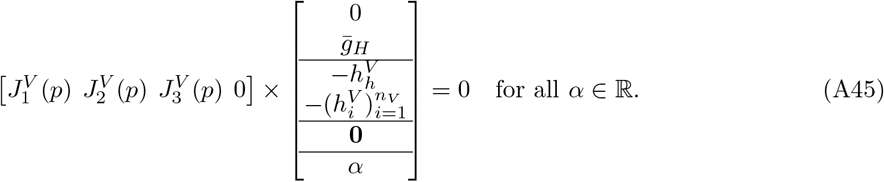 Again, from (A40)–(A43) one can check that the vector in (A45) is in 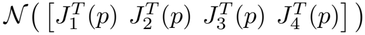 if and only if *α* = 0. Hence, the rank of Jacobian *J*^*V T*^ (*p*) in (A37) is 3 + 2*n*_*V*_ .
ii. Let *n*_*V*_ = 1. By (A35),

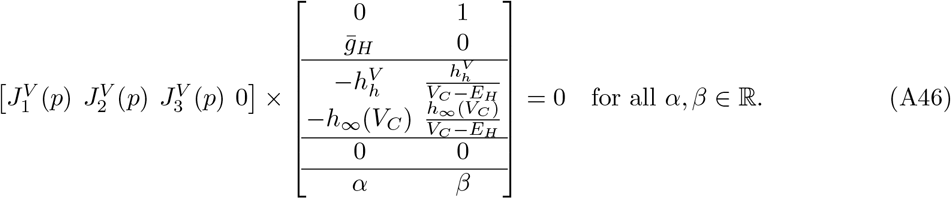

From (A40)–(A43) one can check that the first column vector in (A46) is in 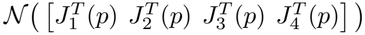 if an d only if *α* = 0, and that the second column vector in (A46) is not in 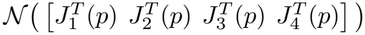 for any *β ∈* ℝ . Hence, the rank of Jacobian *J*^*V T*^ (*p*) in (A37) is 3 + 2*n*_*V*_ .

#### A.6 Proof of Theorem 2.15 (rank *J*^*V*^ (*p*); A-current, whole cell)

This section provides a proof of Theorem 2.15 on the Jacobian rank of the A-current model constrained by the voltage clamp experiment.

We prove the more general result with the power 3 in (2.28) replaced by any *q ∈* ℕ, in which case the condition *m*_*V*_ *≥* 15 in Assumption 2.14 is replaced by *m*_*V*_*≥*4*q* + 3.

We introduce the abbreviations

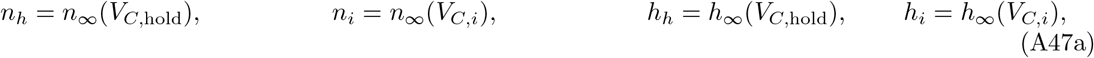

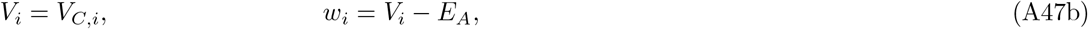

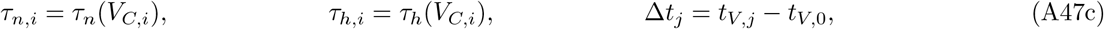

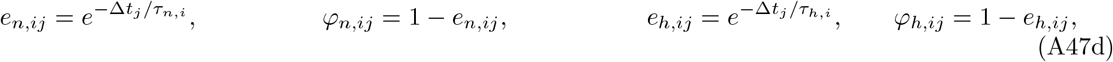

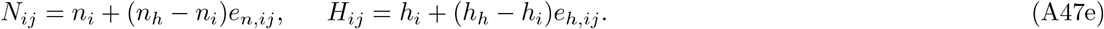

As before, ⊙denotes the entrywise product. Furthermore, we define 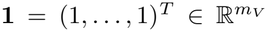, we let **0** = (0, …, 0)^*T*^ denote the zero vector in 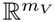 or in 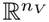, depending on the context, and for each experiment *i* = 1, …, *n*_*V*_, we define the 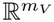 vectors

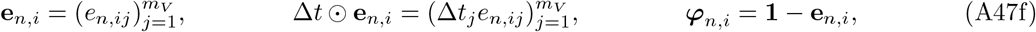

and similar quantities for *h*, and

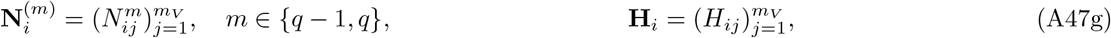

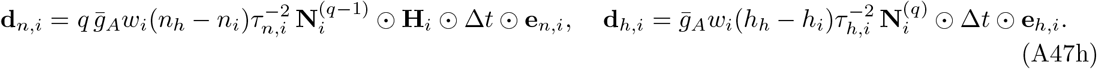

The abbreviations (A47) allow us to rewrite

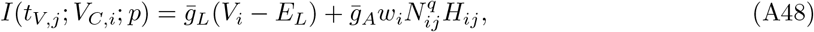

cf. (2.28) with 3 replaced by *q*. The residual 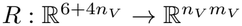,

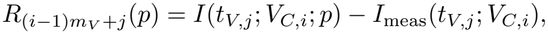

is a block vector with *n*_*V*_ blocks of size *m*_*V*_ .

With 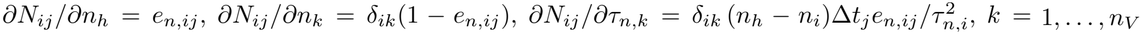, and the analogous formulas for *H*_*ij*_, we get

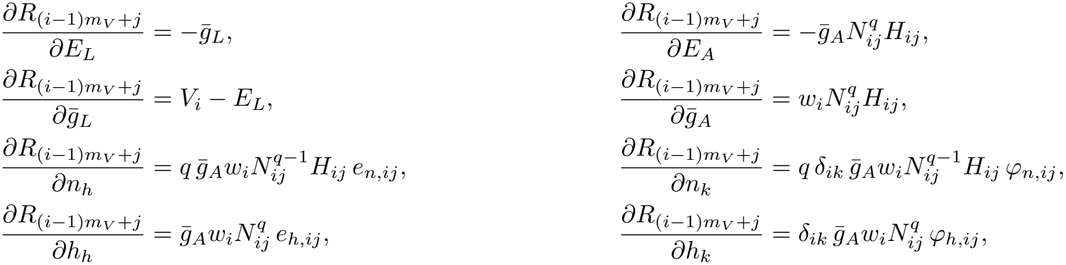

for *k* = 1, …, *n*_*V*_ .

The Jacobian of the residual function corresponding to (2.9) with *I t*_*V,j*_; *V*_*C,i*_; *p* given by the function (2.28) with *V*_*C*_ = *V*_*C,i*_ at times *t*_*V,j*_ *∈* [*t*_*V*,0_, *t*_*V,f*_ ), and columns ordered according to *p* as in (2.29), is given by

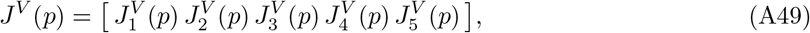

where

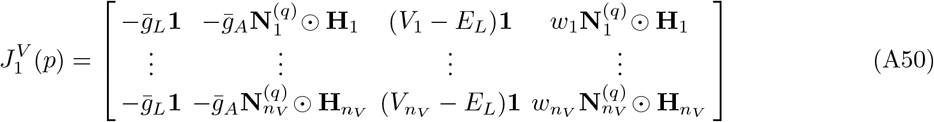

is the partial Jacobian of derivatives with respect to 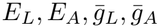,

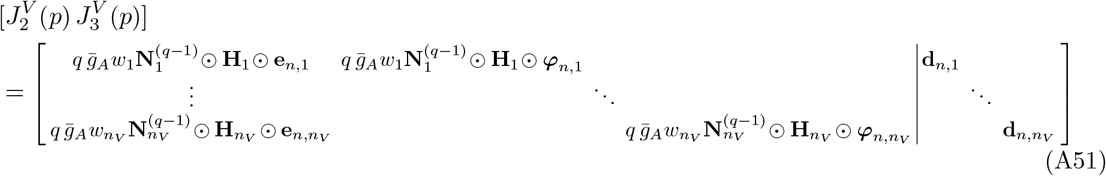

is the partial Jacobian of derivatives with respect to 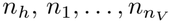 and 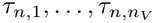, and

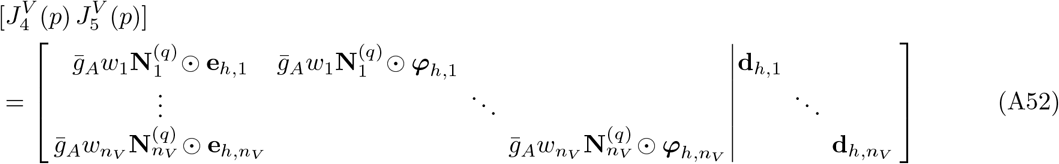

is the partial Jacobian of derivatives with respect to 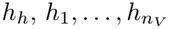 and 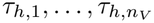.

A direct computation shows that

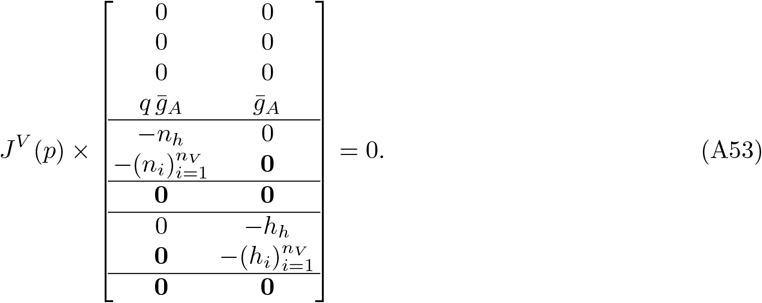

Hence the null space of *J*^*V*^ (*p*) is at least two-dimensional and its rank is at most 4 + 4*n*_*V*_ .

Next, we show that the 4 + 4*n*_*V*_ columns 1, 2, 3, 4, 6, …, 5 + 2*n*_*V*_, 7 + 2*n*_*V*_, …, 6 + 4*n*_*V*_, i.e., all columns except for columns 5 and 6 + 2*n*_*V*_, which correspond to the partial derivatives with respect to *n*_*h*_ and *h*_*h*_, are linearly independent. This implies that the rank of *J*^*V*^ (*p*) is 4 + 4*n*_*V*_ .

We set a linear combination of the 4 + 4*n*_*V*_ columns 1, 2, 3, 4, 6, …, 5 + 2*n*_*V*_, 7 + 2*n*_*V*_, …, 6 + 4*n*_*V*_ to zero and show that the coefficients in this linear combination must be zero. Using (A47), the linear combination reads

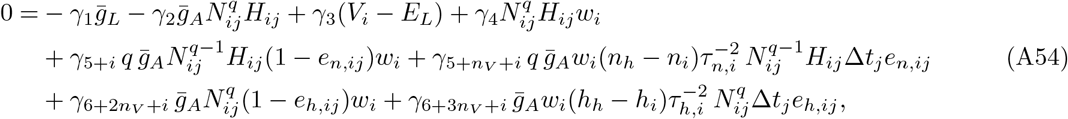

for *j* = 1, …, *m*_*V*_ and *i* = 1, …, *n*_*V*_ . By (A47d), (A47e), *N*_*ij*_, *H*_*ij*_, *e*_*n,ij*_, and *e*_*h,ij*_ are functions of time Δ*t*_*j*_. If we rearrange (A54) according to the different functions of time Δ*t*_*j*_, then (A54) becomes

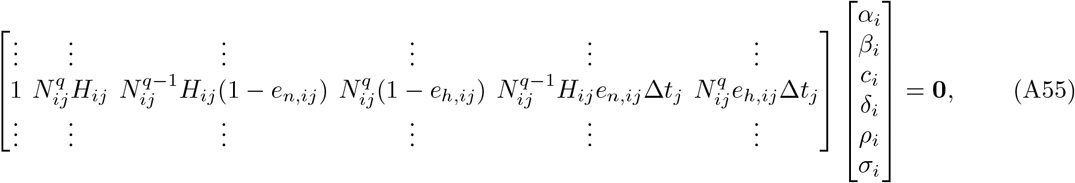

for *i* = 1, …, *n*_*V*_, where

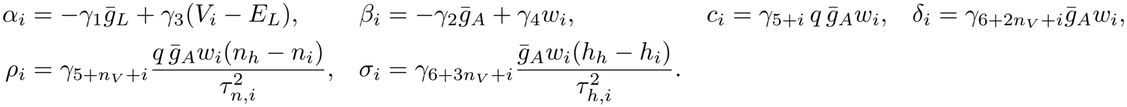

Let 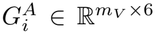 be the matrix in (A55). We will show below that the matrices 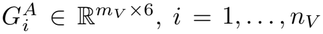, have full column rank. Hence, (A55) implies *α*_*i*_ = *β*_*i*_ = *c*_*i*_ = *δ*_*i*_ = *ρ*_*i*_ = *σ*_*i*_ = 0 for *i* = 1, …, *n*_*V*_ . Since 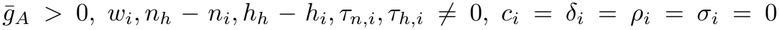 gives 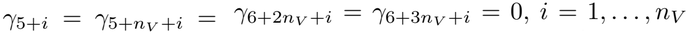. Moreover, 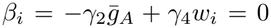 for *i* = 1, …, *n*_*V*_, where *n*_*V*_ *>* 1 and *w*_*i*_ *≠ w*_*k*_ for *i ≠ k* (since *V*_*i*_ *≠ V*_*k*_), implies *γ*_4_ = 0, and then 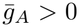 gives *γ*_2_ = 0. Similarly, 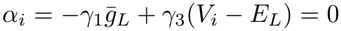 for *i* = 1, …, *n*_*V*_, *n*_*V*_ *>* 1, and *V*_*i*_ *≠ V*_*k*_ for *i ≠ k* imply *γ*_3_ = 0, and then 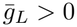 gives *γ*_1_ = 0.

To finish the proof, it remains to show that the matrices 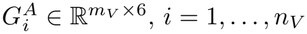, in (A55) have full column rank.

Define the functions

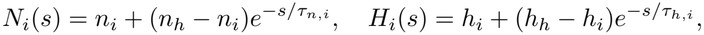

and

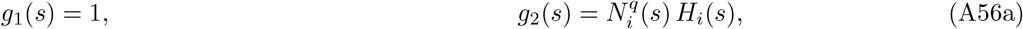

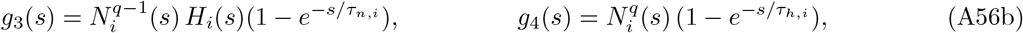

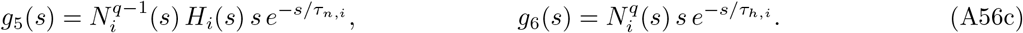

The columns of the matrix 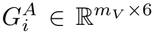 in (A55) are the functions *g*_1_, …, *g*_6_ evaluated at *s* = Δ*t*_*j*_, *j* = 1, …, *m*_*V*_ . The full rank property of 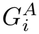 is stated in Lemma A.3 below.

##### Lemma A.1.

*Let q ∈* ℕ. *If τ*_*n,i*_, *≠ τ*_*h,i*_ *>* 0 *with τ*_*n,i*_ ≠ *τ*_*h,i*_, *n*_*h*_ 0, *h*_*h*_ ≠ 0, *n*_*h*_ *≠ n*_*i*_, *h*_*h*_ *≠ h*_*i*_, *and* (*n*_*i*_, *h*_*i*_) *≠* (0, 0), *then the functions g*_1_, …, *g*_6_ *in* (A56) *are linearly independent as functions of s ∈* ℝ.

*Proof* We write *λ* = 1*/τ*_*n,i*_, *µ* = 1*/τ*_*h,i*_, *E*_*n*_(*s*) = *e*^*−λs*^, *E*_*h*_(*s*) = *e*^*−µs*^, so that *N*_*i*_(*s*) = *n*_*i*_ + (*n*_*h*_ *− n*_*i*_)*E*_*n*_(*s*), *H*_*i*_(*s*) = *h*_*i*_ + (*h*_*h*_ *− h*_*i*_)*E*_*h*_(*s*).

Expanding the powers of *N*_*i*_, every *g*_*k*_ is a finite linear combination of functions *s* ^*m*^*e*^*−νs*^, *m ∈ {*0, 1*}*, with exponents *ν* = *kλ* + *lµ ≥* 0, *k ∈ {*0, …, *q}, l ∈ {*0, 1*}*. We use that the functions *s*^*m*^*e*^*−νs*^ with distinct pairs (*m, ν*) are linearly independent on ℝ.

Suppose 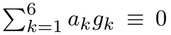. Since *g*_1_, …, *g*_4_ contain only terms *s*^*m*^*e*^*−νs*^ with *m* = 0 and *g*_5_, *g*_6_ only terms *s*^*m*^*e*^*−νs*^ with *m* = 1, both groups vanish separately:

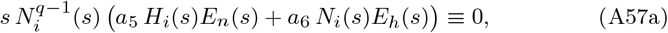

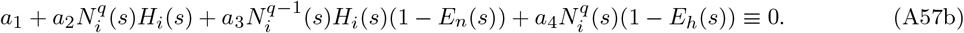

First we prove *a*_5_ = *a*_6_ = 0. Since *s ⟼ N*_*i*_(*s*) = *n*_*i*_ + (*n*_*h*_ *− n*_*i*_)*e*^*−λs*^ is strictly monotone, the factor 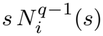 vanishes at most at two points. By continuity, (A57a) implies

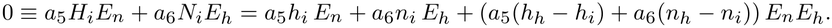

Since *λ ≠ µ*, the exponents *λ, µ, λ* + *µ* are distinct, so *a*_5_*h*_*i*_ = 0, *a*_6_*n*_*i*_ = 0, and *a*_5_(*h*_*h*_ *− h*_*i*_) + *a*_6_(*n*_*h*_ *− n*_*i*_) = 0. If *h*_*i*_ *≠* 0, then *a*_5_ = 0, hence *a*_6_(*n*_*h*_ *− n*_*i*_) = 0 and *a*_6_ = 0. If *n*_*i*_ ≠ 0, then *a*_6_ = 0, hence *a*_5_(*h*_*h*_ *− h*_*i*_) = 0 and *a*_5_ = 0.

Next, we prove *a*_1_ = *· · ·* = *a*_4_ = 0. Inserting *H*_*i*_ = *h*_*i*_ + (*h*_*h*_ *− h*_*i*_)*E*_*h*_ into (A57b) gives

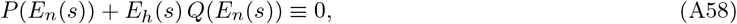

with the polynomials (in *x* = *E*_*n*_(*s*)) *P* and *Q* given by

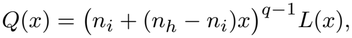

with

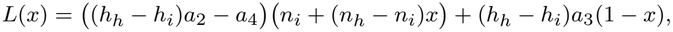

and

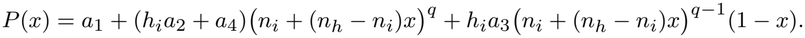

Both *P* and *Q* are polynomials of degree at most *q*, i.e., deg *P*, deg *Q ≤ q*.

The exponents occurring in *P* (*E*_*n*_) lie in *{kλ* : 0 *≤ k ≤ q}*, those in *E*_*h*_*Q*(*E*_*n*_) lie in *{µ* + *kλ* : 0 *≤ k ≤ q}*.

These two sets intersect only if *µ* = *mλ* for an integer *m ∈ {*2, …, *q}* (recall *µ >* 0 and *µ ≠ λ*).

- If the sets are disjoint, then (A58) implies *E*_*h*_*Q*(*E*_*n*_) *≡* 0. Thus, *Q* vanishes on the interval (0, *∞*), the range of *E*_*n*_, which implies *Q ≡* 0.
- If *µ* = *mλ* with *m ∈ {*2, …, *q}*, then 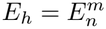 and (A58) implies *P* (*x*) + *x*^*m*^*Q*(*x*) = 0 for *x ∈* (0, *∞*). If *Q ≠* 0, then *L ≠* 0 and deg *Q ≥ q −* 1, so that deg(*x*^*m*^*Q*) *≥ m* + *q −* 1 *≥ q* + 1 *>* deg *P*, a contradiction to deg *P ≤ q*. Hence *Q ≡* 0.

Since (*n*_*h*_ *− n*_*i*_) ≠0, (*n*_*i*_ + (*n*_*h*_ *− n*_*i*_)*x*) is a nonzero polynomial, the identity *Q ≡* 0 implies *L ≡* 0, i.e.,

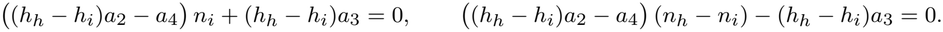

Adding both equations gives (*h*_*h*_ *− h*_*i*_)*a*_2_ *− a*_4_ *n*_*i*_ + (*n*_*h*_ *− n*_*i*_) = (*h*_*h*_ *− h*_*i*_)*a*_2_ *− a*_4_ *n*_*h*_ = 0, and *n*_*h*_ *≠* 0 yields *a*_4_ = (*h*_*h*_ *− h*_*i*_)*a*_2_. Then (*h*_*h*_ *− h*_*i*_)*a*_3_ = 0, and *h*_*h*_ *− h*_*i*_ ≠ 0 gives *a*_3_ = 0.

Next, *Q ≡* 0 and (A58) give *P* (*E*_*n*_) *≡* 0. Since *a*_3_ = 0, *a*_4_ = (*h*_*h*_ *− h*_*i*_)*a*_2_, and *E*_*n*_ takes all values in (0, *∞*),

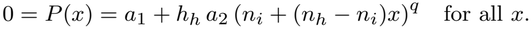

Since *n*_*h*_ *− n*_*i*_ ≠0 and *q ≥* 1, *n*_*i*_ + (*n*_*h*_ *− n*_*i*_)*x* ^*q*^ is a nonconstant polynomial, so *a*_1_ = 0 and *h*_*h*_*a*_2_ = 0. Finally, *h*_*h*_ *≠* 0 gives *a*_2_ = 0 and *a*_4_ = (*h*_*h*_ *− h*_*i*_)*a*_2_ = 0.

Note that the assumptions *h*_*h*_ *≠*0, *n*_*h*_ *≠* 0, *n*_*h*_ *≠ n*_*i*_, and (*n*_*i*_, *h*_*i*_) ≠ (0, 0) of Lemma A.1 cannot be dropped: If *h*_*h*_ = 0, then *g*_2_ = *h*_*i*_ *g*_4_; if *n*_*h*_ = 0, then *g*_2_ = *n*_*i*_ *g*_3_; if *n*_*h*_ = *n*_*i*_, then 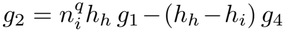 and if *n*_*i*_ = *h*_*i*_ = 0, then *g*_5_ and *g*_6_ are both multiples of *s* 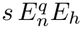.

The next result is needed in the proof of Lemma A.3 below to conclude that the linear independence of the functions *g*_1_, …, *g*_6_ in (A56) implies linear independence of the columns of the matrix 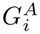 in (A55), which consist of values of *g*_1_, …, *g*_6_ at times 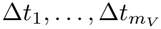.

##### Lemma A.2.

*Let µ*_1_, …, *µ*_*K*_ *be real numbers (not necessarily distinct). Every nonzero real solution f of the linear ODE*

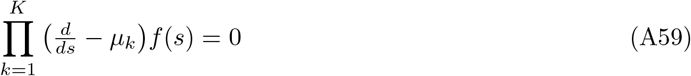

*has at most K −* 1 *distinct real zeros*.

*Proof* The proof is by induction on *K*. For 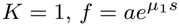 with *a ≠* 0 has no zeros.

Let the statement hold for *K*. We prove the induction step by contradiction. Let *f* be a nonzero solution of 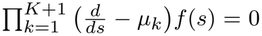 with at least *K* + 1 distinct zeros. Then 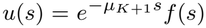 has the same zeros, and by Rolle’s theorem *u*^*′*^ has at least *K* distinct zeros, and so does 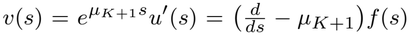. Since the operators 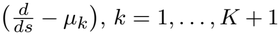, commute, *v* solves the ODE (A59) with the *K* factors *µ*_1_, …, *µ*_*K*_ . By induction *v ≡* 0, i.e., 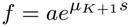, which has a zero only if *a* = 0, contradicting *f ≠* 0.

##### Lemma A.3.

*Under the assumptions of Lemma A*.*1, if the times* 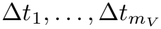*contain at least* 4*q* + 3 *distinct values, then* 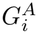*in* (A55) *has rank* 6.

*Proof* Expanding the powers of *N*_*i*_, every function *g*_*k*_ in (A56) is a linear combination of the functions 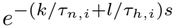 with *k ∈ {*0, …, *q}, l ∈ {*0, 1*}* (at most 2*q* + 2 distinct exponents), and of the functions 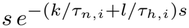 with *k ∈ {*0, …, *q}, l ∈ {*0, 1*}*, (*k, l*) *≠* (0, 0) (at most 2*q* + 1 distinct exponents). Hence every linear combination of *g*_1_, …, *g*_6_ solves a linear constant-coefficient ODE of the form (A59) of order at most (2*q* + 2) + (2*q* + 1) = 4*q* + 3 with real characteristic roots (Teschl 2012, Sec. 3.3).

By Lemma A.1, a nonzero coefficient vector gives a nonzero solution, which by Lemma A.2 has at most 4*q* + 2 distinct zeros and therefore cannot vanish at 4*q* + 3 distinct sample times.

For *q* = 3, it suffices that the times 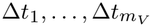 contain at least 15 distinct values.

#### A.7 Proof of Theorem 2.18 (rank *J*^*V*^ (*p*); K-current, whole cell)

This section provides a proof of Theorem 2.18 on the Jacobian rank of the delayed rectifier potassium current model constrained by the voltage clamp experiment.

We prove the more general result with the power 4 in (2.32) replaced by any integer *q ≥* 2, in which case the condition *m*_*V*_ *≥* 9 in Assumption 2.17 is replaced by *m*_*V*_ *≥* 2*q* + 1. (For *q* = 1, see the remark after Lemma A.4.)

We introduce the abbreviations

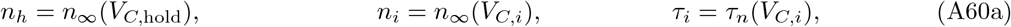

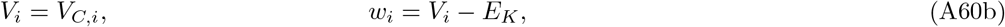

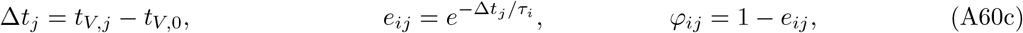

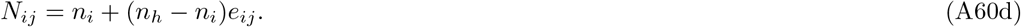

As before, *⊙* denotes the entrywise product and 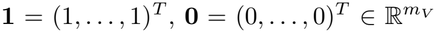. Furthermore, for each experiment *i* = 1, …, *n*_*V*_, we define the 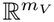 vectors

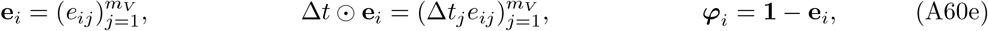

and

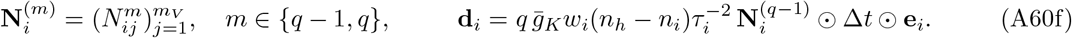

With the notation (A60), the current (2.32) at *V*_*C*_ = *V*_*C,i*_ and *t* = *t*_*V,j*_ is written as

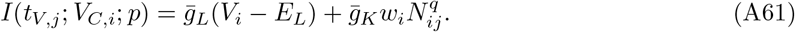

The residual 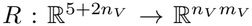 is a vector of *n*_*V*_ blocks, each of length *m*_*V*_, and has components given by (A4). Using *∂N*_*ij*_*/∂n*_*h*_ = *e*_*ij*_, *∂N*_*ij*_*/∂n*_*k*_ = *δ*_*ik*_*φ*_*ij*_ and 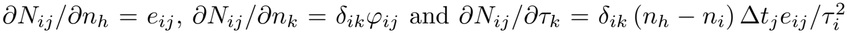, the partial derivatives of the residual are

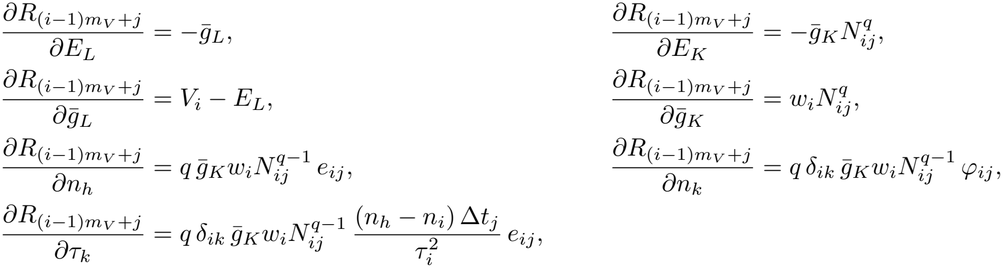

for *k* = 1, …, *n*_*V*_ .

Using the column order of *p* as in (2.33), 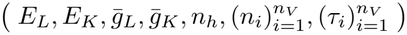, the Jacobian in block form is

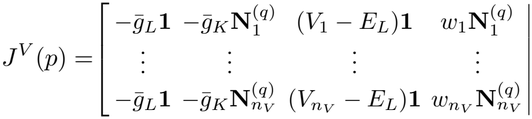

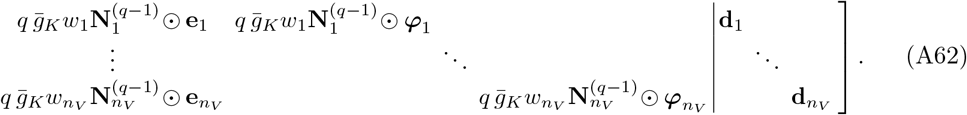

Defining

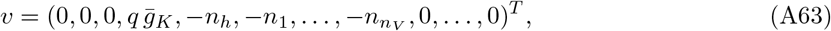

a direct computation shows that

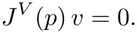

Hence the null space of *J*^*V*^ (*p*) is at least one-dimensional and its rank is at most 4 + 2*n*_*V*_ . Next, we show that the 4 + 2*n*_*V*_ columns 1, 2, 3, 4, 5 + *i, i* = 1, …, 2*n*_*V*_, i.e., all columns except column 5, which corresponds to the partial derivative with respect to *n*_*h*_, are linearly independent. This implies that the rank of *J*^*V*^ (*p*) is 4 + 2*n*_*V*_ .

To show that the columns 1, 2, 3, 4, 5 + *i, i* = 1, …, 2*n*_*V*_, are linearly independent, we set a linear combination of them to zero and show that the coefficients must be zero. Let

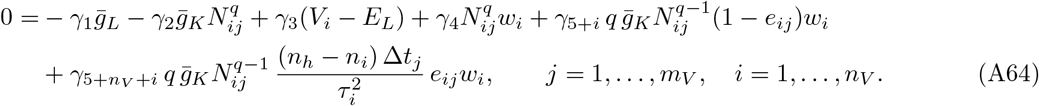

By (A60), *N*_*ij*_ and *e*_*ij*_ are functions of time Δ*t*_*j*_. If we rearrange (A64) according to the different functions of time Δ*t*_*j*_, then (A64) becomes

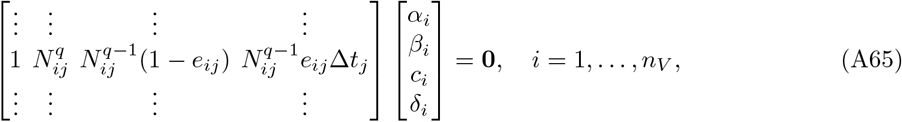

with

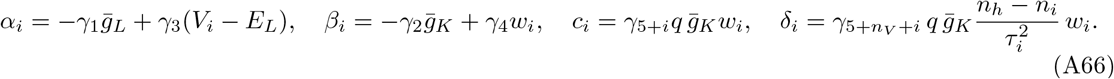

Let 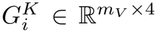 be the matrix in (A65). We will show below that the matrices 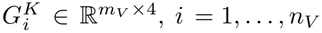, have full column rank. Hence, (A65) implies *α*_*i*_ = *β*_*i*_ = *c*_*i*_ = *δ*_*i*_ = 0 for *i* = 1, …, *n*_*V*_ . Since 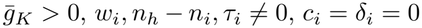 gives *γ*_5+*i*_ = 0, *i* = 1, …, 2*n*_*V*_ . Moreover, 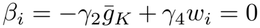 for *i* = 1, …, *n*_*V*_, where *n*_*V*_ *>* 1 and *w*_*i*_ *≠ w*_*k*_ for *i ≠ k* (since *V*_*i*_ *≠ V*_*k*_), implies *γ*_4_ = 0, and then 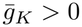 gives *γ*_2_ = 0. Similarly, 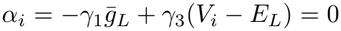 for *i* = 1, …, *n*_*V*_, *n*_*V*_ *>* 1, and *V*_*i*_ ≠ *V*_*k*_ for *i ≠ k* imply *γ*_3_ = 0, and then 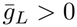 gives *γ*_1_ = 0.

To finish the proof, it remains to show that the matrices 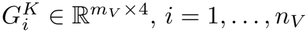, in (A65) have full column rank.

Define the functions

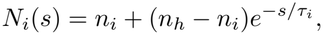

and

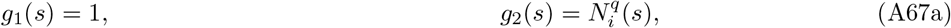

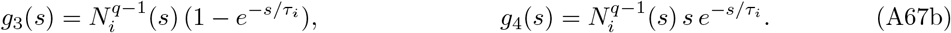

The columns of the matrix 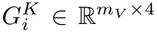 in (A65) are the functions *g*_1_, …, *g*_4_ evaluated at *s* = Δ*t*_*j*_, *j* = 1, …, *m*_*V*_ . The full rank property of 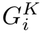 is established in Lemma A.5 below.

##### Lemma A.4.

*Let q ∈* ℕ *with q ≥* 2, *τ*_*i*_ *>* 0, *n*_*h*_ *n*_*i*_. *Then the functions g*_1_, …, *g*_4_ *in* (A67) *are linearly independent as functions of s ∈* ℝ.

*Proof* We write *λ* = 1*/τ*_*i*_ and *E*(*s*) = *e*^*−λs*^ so that *N*_*i*_(*s*) = *n*_*i*_ + (*n*_*h*_ *− n*_*i*_)*E*(*s*). Expanding the powers of *N*_*i*_, the functions *g*_1_, *g*_2_, *g*_3_ are linear combinations of *e*^*−kλs*^, *k ∈ {*0, …, *q}*, and *g*_4_ is a linear combination of *s e*^*−kλs*^, *k ∈ {*1, …, *q}*. Since functions *s*^*m*^*e*^*−νs*^ with distinct pairs (*m, ν*) are linearly independent on 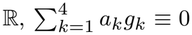 implies

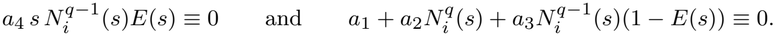

Since 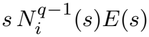 vanishes at most at two points, the first identity gives *a*_4_ = 0. Since *E*(*s*) *∈* (0, *∞*), we view the second identity as an identity in *x* = *E*(*s*). Note that because the resulting identity in *x* is a polynomial identity, it holds for all *x ∈* ℝ. Hence, we have the polynomial identity

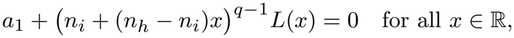

where

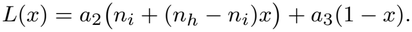

Evaluating at *x* = *−n*_*i*_*/*(*n*_*h*_*−n*_*i*_) and using *q ≥* 2 gives *a*_1_ = 0. Since (*n*_*i*_+(*n*_*h*_*−n*_*i*_)*x*)^*q−*1^ *≠* 0 for *x ≠ −n*_*i*_*/*(*n*_*h*_*−n*_*i*_), and *L* is continuous, *L*(*x*) = 0, *x ∈* ℝ, i.e., *a*_2_*n*_*i*_ + *a*_3_ = 0 and *a*_2_(*n*_*h*_ *− n*_*i*_) *− a*_3_ = 0. Adding both equations gives *a*_2_*n*_*h*_ = 0, so *a*_2_ = 0 by *n*_*h*_ *≠* 0, and then *a*_3_ = 0.

The assumptions of Lemma A.4 cannot be dropped: if *n*_*h*_ = 0, then *g*_2_ = *n*_*i*_ *g*_3_; if *n*_*h*_ = *n*_*i*_, then 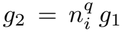 and if *q* = 1, then *g*_2_ = *n*_*h*_ *g*_1_ *−* (*n*_*h*_ *− n*_*i*_) *g*_3_. For *q* = 1 the situation is the same as in Section 2.2.1: Theorem 2.2 applies, and the rank deficiency of *J*^*V*^ (*p*) is three.

##### Lemma A.5.

*Under the assumptions of Lemma A*.*4, if the times* 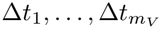*contain at least* 2*q* + 1 *distinct values, then* 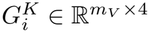 *in* (A65) *has rank* 4.

*Proof* Every function *g*_1_, …, *g*_4_ in (A67) is a linear combination of the functions 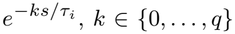 (*q* + 1 exponents), and 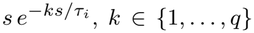 (*q* exponents). Hence every linear combination of *g*_1_, …, *g*_4_ solves a linear constant-coefficient ODE of the form (A59) of order (*q* + 1) + *q* = 2*q* + 1 with real characteristic roots 0 (simple) and *−k/τ*_*i*_, *k* = 1, …, *q* (each double) (Teschl 2012, Sec. 3.3). By Lemma A.4, a nonzero coefficient vector gives a nonzero solution, which by Lemma A.2 in Section A.6 has at most 2*q* distinct zeros and therefore cannot vanish at 2*q* + 1 distinct sample times.

For *q* = 4, it suffices that the times 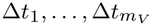 contain at least 9 distinct values.

## References

Achard P, De Schutter E (2006) Complex Parameter Landscape for a Complex Neuron Model. PLOS Computational Biology 2(7):e94. 10.1371/journal.pcbi.0020094

Angelo K, London M, Christensen SR, et al (2007) Local and global effects of Ih distribution in dendrites of mammalian neurons. Journal of Neuroscience 27(32):8643–8653. 10.1523/JNEUROSCI.5284-06.2007

Bard Y (1974) Nonlinear Parameter Estimation. Academic Press [Harcourt Brace Jovanovich, Publishers], New York-London

Björck Å (2024) Numerical Methods for Least Squares Problems, 2nd edn. Society for Industrial and Applied Mathematics (SIAM), Philadelphia, PA, 10.1137/1.9781611977950

Csercsik D, Hangos KM, Szederkényi G (2012) Identifiability analysis and parameter estimation of a single Hodgkin-Huxley type voltage dependent ion channel under voltage step measurement conditions. Neurocomputing 77:178–188. 10.1016/j.neucom.2011.09.006

Deistler M, Kadhim KL, Pals M, et al (2025) Jaxley: differentiable simulation enables large-scale training of detailed biophysical models of neural dynamics. Nature Methods 22(12):2649–2657. 10.1038/s41592-025-02895-w

Dennis JE Jr., Schnabel RB (1996) Numerical Methods for Unconstrained Optimization and Nonlinear Equations, Classics in Applied Mathematics, vol 16. Society for Industrial and Applied Mathematics (SIAM), Philadelphia, PA, 10.1137/1.9781611971200, corrected reprint of the 1983 original

Donaldson JR, Schnabel RB (1987) Computational experience with confidence regions and confidence intervals for nonlinear least squares. Technometrics 29(1):67–82. 10.2307/1269884

Druckmann S, Banitt Y, Gidon A, et al (2007) A novel multiple objective optimization framework for constraining conductance-based neuron models by experimental data. Frontiers in Neuroscience 1(1):7–18. 10.3389/neuro.01.1.1.001.2007

Eriksson J, Wedin PA, Gulliksson ME, et al (2005) Regularization methods for uniformly rank-deficient nonlinear least-squares problems. J Optim Theory Appl 127(1):1–26. 10.1007/s10957-005-6389-0

Fracasso K, Heinkenschloss M (2027) Solution manifolds for nonlinear least squares problems, in preparation

Goaillard JM, Marder E (2021) Ion channel degeneracy, variability, and covariation in neuron and circuit resilience. Annual Review of Neuroscience 44:335–357. 10.1146/annurev-neuro-092920-121538

Golowasch J, Marder E (1992) Ionic currents of the lateral pyloric neuron of the stomatogastric ganglion of the crab. Journal of Neurophysiology 67(2):318–331. 10.1152/jn.1992.67.2.318

Hamill OP, Marty A, Neher E, et al (1981) Improved patch-clamp techniques for high-resolution current recording from cells and cell-free membrane patches. Pflügers Archiv 391(2):85–100. 10.1007/BF00656997

Hansen PC (1998) Rank-Deficient and Discrete Ill-Posed Problems. Numerical Aspects of Linear Inversion. SIAM Monographs on Mathematical Modeling and Computation, Society for Industrial and Applied Mathematics (SIAM), Philadelphia, PA, 10.1137/1.9780898719697

He C, Chen F, Li B, et al (2014) Neurophysiology of HCN channels: From cellular functions to multiple regulations. Progress in Neurobiology 112:1–23. 10.1016/j.pneurobio.2013.10.001

Hille B (2001) Ion Channels of Excitable Membranes, 3rd edn. Oxford University Press, Oxford, UK

Hodgkin AL, Huxley AF (1952) A quantitative description of membrane current and its application to conduction and excitation in nerve. The Journal of Physiology 117(4):500–544. 10.1113/jphysiol.1952.sp004764

Huys QJM, Ahrens MB, Paninski L (2006) Efficient estimation of detailed single-neuron models. Journal of Neurophysiology 96(2):872–890. 10.1152/jn.00079.2006

Keren N, Peled N, Korngreen A (2005) Constraining compartmental models using multiple voltage recordings and genetic algorithms. Journal of Neurophysiology 94(6):3730–3742. 10.1152/jn.00408.2005

Liu Z, Golowasch J, Marder E, et al (1998) A model neuron with activity-dependent conductances regulated by multiple calcium sensors. Journal of Neuroscience 18(7):2309–2320. 10.1523/JNEUROSCI.18-07-02309.1998

Magee JC (1998) Dendritic hyperpolarization-activated currents modify the integrative properties of hippocampal CA1 pyramidal neurons. Journal of Neuroscience 18(19):7613–7624. 10.1523/JNEUROSCI.18-19-07613.1998

Namazifard S, Gabbiani F, Heinkenschloss M (2026) A toolkit to identify parameter degeneracy and solution manifolds of Hodgkin-Huxley type current models. bioRxiv preprint

Nocedal J, Wright SJ (2006) Numerical Optimization, 2nd edn. Springer Verlag, Berlin, Heidelberg, New York, 10.1007/978-0-387-40065-5

Prinz AA, Thirumalai V, Marder E (2003) The functional consequences of changes in the strength and duration of synaptic inputs to oscillatory neurons. Journal of Neuroscience 23(3):943–954. 10.1523/JNEUROSCI.23-03-00943.2003

Prinz AA, Bucher D, Marder E (2004) Similar network activity from disparate circuit parameters. Nature Neuroscience 7(12):1345–1352. 10.1038/nn1352

Rall W (1962) Electrophysiology of a dendritic neuron model. Biophysical Journal 2(2 Pt 2):145–167. 10.1016/S0006-3495(62)86953-7

Seber GAF, Wild CJ (2003) Nonlinear Regression. Wiley-Interscience Paperback Series, John Wiley & Sons, Inc., New York, 10.1002/0471725315

Teschl G (2012) Ordinary differential equations and dynamical systems, Graduate Studies in Mathematics, vol 140. American Mathematical Society, Providence, RI, 10.1090/gsm/140

Walch OJ, Eisenberg MC (2016) Parameter identifiability and identifiable combinations in generalized Hodgkin-Huxley models. Neurocomputing 199:137–143. 10.1016/j.neucom.2016.03.027

Willms AR, Baro DJ, Harris-Warrick RM, et al (1999) An improved parameter estimation method for Hodgkin-Huxley models. J Comput Neurosci 6:145–168. 10.1023/A:1008880518515

